# Spatial proteomics in neurons at single-protein resolution

**DOI:** 10.1101/2023.05.17.541210

**Authors:** Eduard M. Unterauer, Sayedali Shetab Boushehri, Kristina Jevdokimenko, Luciano A. Masullo, Mahipal Ganji, Shama Sograte-Idrissi, Rafal Kowalewski, Sebastian Strauss, Susanne C.M. Reinhardt, Ana Perovic, Carsten Marr, Felipe Opazo, Eugenio F. Fornasiero, Ralf Jungmann

**Affiliations:** Max Planck Institute of Biochemistry, Planegg, Germany; Faculty of Physics and Center for NanoScience, Ludwig-Maximilians-Universität, Munich, Germany; Institute of AI for Health, Helmholtz Zentrum München - German Research Center for Environmental Health, Neuherberg, Germany; Data & Analytics, Pharmaceutical Research and Early Development, Roche Innovation Center Munich (RICM), Penzberg, Germany; Department of Mathematics, Technical University of Munich, Munich, Germany; Institute of Neuro- and Sensory Physiology, University Medical Center Göttingen, Göttingen, Germany; Center for Biostructural Imaging of Neurodegeneration (BIN), University Medical Center Göttingen, Göttingen, Germany; NanoTag Biotechnologies GmbH, Göttingen, Germany; Department of Life Sciences, University of Trieste, Trieste, Italy; These authors contributed equally; Lead contact

## Abstract

To fully understand biological processes and functions, it is necessary to reveal the molecular heterogeneity of cells and even subcellular assemblies by gaining access to the location and interaction of all biomolecules. The study of protein arrangements has seen significant advancements through super-resolution microscopy, but such methods are still far from reaching the multiplexing capacity of spatial proteomics. Here, we introduce Secondary label-based Unlimited Multiplexed DNA-PAINT (SUM-PAINT), a high-throughput imaging method capable of achieving virtually unlimited multiplexing at better than 15 nm spatial resolution. Using SUM-PAINT, we generated the most extensive multiprotein dataset to date at single-protein spatial resolution, comprising up to 30 distinct protein targets in parallel and adapted omics-inspired analysis workflows to explore these feature-rich datasets. Remarkably, by examining the multiplexed protein content of almost 900 individual synapses at single-protein resolution, we revealed the complexity of synaptic heterogeneity, ultimately leading to the discovery of a new synapse type. This work provides not only a feature-rich resource for researchers, but also an integrated data acquisition and analysis workflow for comprehensive spatial proteomics at single-protein resolution, paving the way for ‘Localizomics’.

## INTRODUCTION

One of the key goals of life science research is to understand living systems from the level of whole organisms and tissues to the assembly of complex cellular networks and eventually the organization and interaction of individual biomolecules. To achieve this goal, we need technologies that can accurately measure all biomolecules and their spatial interactions in a truly quantitative manner. Omics techniques are poised to meet this need. Traditional omics-based technologies such as sequencing or mass spectrometry have a profound impact on life science research by quantifying nucleic acid and protein abundance even in complex tissue environments. However, to fully understand biological processes and functions, it is necessary to reveal the molecular heterogeneity of cells and even subcellular assemblies by gaining access to the location and interaction of all biomolecules. This goal has been the focus of spatial omics techniques developed in recent years, which provide both absolute quantification and subcellular localization of molecular assemblies.

Methods like MERFISH^1^ and seqFISH+^2^ have been developed to offer spatially resolved transcriptomics and genomics, recently contributing to the assembly of cell atlases depicting age-related changes in mouse frontal cortices^3^, as well as characterizing the chromatin states and nuclear organization within specific cell types^4^. However, while these approaches to subcellular DNA and RNA imaging are pushing the boundaries of spatially-resolved single-cell biology, a comparable technology for mapping single proteins is still missing. To achieve a comprehensive understanding of protein organization at the nanoscale, four critical challenges need to be addressed: sensitivity, throughput, spatial resolution, and multiplexing capabilities.

Mass spectrometry-based proteomics is the leading method for protein quantification, offering exceptional multiplexing and throughput capabilities. To incorporate spatial information, cutting-edge techniques have integrated approaches such as sample dissection^5^, which currently achieve single-cell resolution for up to 5000 different proteins. For attaining subcellular resolution, the employment of specialized labeling reagents has emerged as a highly effective tool. Techniques like imaging mass spectrometry^6^, multiplexed ion beam imaging^7^ and CODEX^8^ utilize the specific labeling of antibodies conjugated with metal ions or DNA, enabling subcellular profiling of tissue samples with up to 100 targets and a spatial resolution of up to 260 nm.

However, the size of most proteins lies in the 5-10 nm range, approximately two orders of magnitude below such a resolution limit. The investigation of protein arrangements has seen significant advancements through super-resolution microscopy^9–11^. Nonetheless, many of these techniques suffer from considerable limitations when it comes to highly multiplexed imaging.

DNA-PAINT is a super-resolution technique that relies on the transient binding of dye-labeled "imager" strands to their complementary "docking" strands present on target molecules of interest^12, 13^. This approach facilitates conceptually unlimited multiplexing through DNA-barcoded sequential imaging, also known as Exchange-PAINT^14^. However, the throughput of DNA-PAINT has traditionally been constrained by its relatively slow image acquisition process. Using optimized *de novo* sequence design and concatenated, repetitive sequence motifs, the acquisition speed of DNA-PAINT has recently been increased^15, 16^ by up to a factor of 100. However, this sequence optimization significantly reduces its multiplexing capacity, effectively limiting it to only six targets.

Here, we introduce Secondary label-based Unlimited Multiplexed PAINT (SUM-PAINT), a method capable of achieving unlimited multiplexing while maintaining the throughput improvement offered by optimized sequence design. We achieved this by decoupling the DNA barcoding of the target from the imaging process using a primary barcode and secondary label.

We demonstrated this technique by generating hippocampal neuron atlases through imaging up to 30 protein targets at single-protein resolution. These unprecedented neuronal architecture maps allowed us not only to resolve spectrin rings, clathrin pits, and synapses with highest spatial resolution, but also to uncover the nano-arrangement of neurofilament fibers and to demonstrate that individual trans-synaptic nanocolumns exhibit precise molecular matches at the single-protein level.

The unprecedented spatial resolution and multiplexing levels of our super-resolution atlases allowed us now for the first time to adapt advanced approaches from machine learning and omics-inspired analyses workflows to explore these feature-rich datasets. Remarkably, by examining the multiplexed protein content of almost 900 individual synapses at single-protein resolution, we uncovered distinctive protein expression patterns and arrangements that truly reveal the synaptic heterogeneity of both excitatory and inhibitory synapses.

This integrated AI-guided analysis workflow, combined with the ability to study each individual synapse down to individual proteins, allowed us to identify a thus far unreported third synapse class. This novel class is defined by the juxtaposition of an inhibitory synapse’s postsynaptic scaffold combined with an excitatory vesicle pool.

### Secondary-label-based DNA-PAINT enables highly-multiplexed imaging of single proteins

Essential requirements for achieving spatial proteomics at the single-protein level using super-resolution microscopy are high spatial resolution and multiplexing capacity at adequate throughput, which is paramount for sufficient statistics. A method that combines all these capabilities has not been established to date.

While DNA-barcoded multiplexing approaches such as Exchange-PAINT^14^ theoretically enable unlimited levels of multiplexing, their throughput has been limited by challenging sequence design, leading to reduced association kinetics. Recent advancements in speed-optimized DNA-PAINT^15, 16^ have led to a 100-fold improvement in throughput by using repetitive, hairpin-free, and concatenated sequence motifs. However, these requirements limit the number of orthogonal sequence motifs to six, and thus multiplexing to only six targets.

To overcome this limitation, we devised a strategy to decouple the speed-optimized target imaging from the sequential multiplexing process using a primary barcode and a secondary label in an approach called Secondary label-based Unlimited Multiplexed (SUM)-PAINT. SUM-PAINT relies on DNA-PAINT imaging of a secondary label (Figure 1A). In this workflow, a primary target of interest is conjugated to a unique 20-nt long DNA sequence: the primary barcode. In a second step, a secondary label, containing a 20-nt complement to the primary barcode, is introduced. This secondary label also contains a speed-optimized docking sequence and a 10-nt long toehold for signal extinction via toehold-mediated strand displacement^17^. In a third step, dye-labeled, speed-optimized imager strands are added to visualize the position of the target.

**Figure 1.**
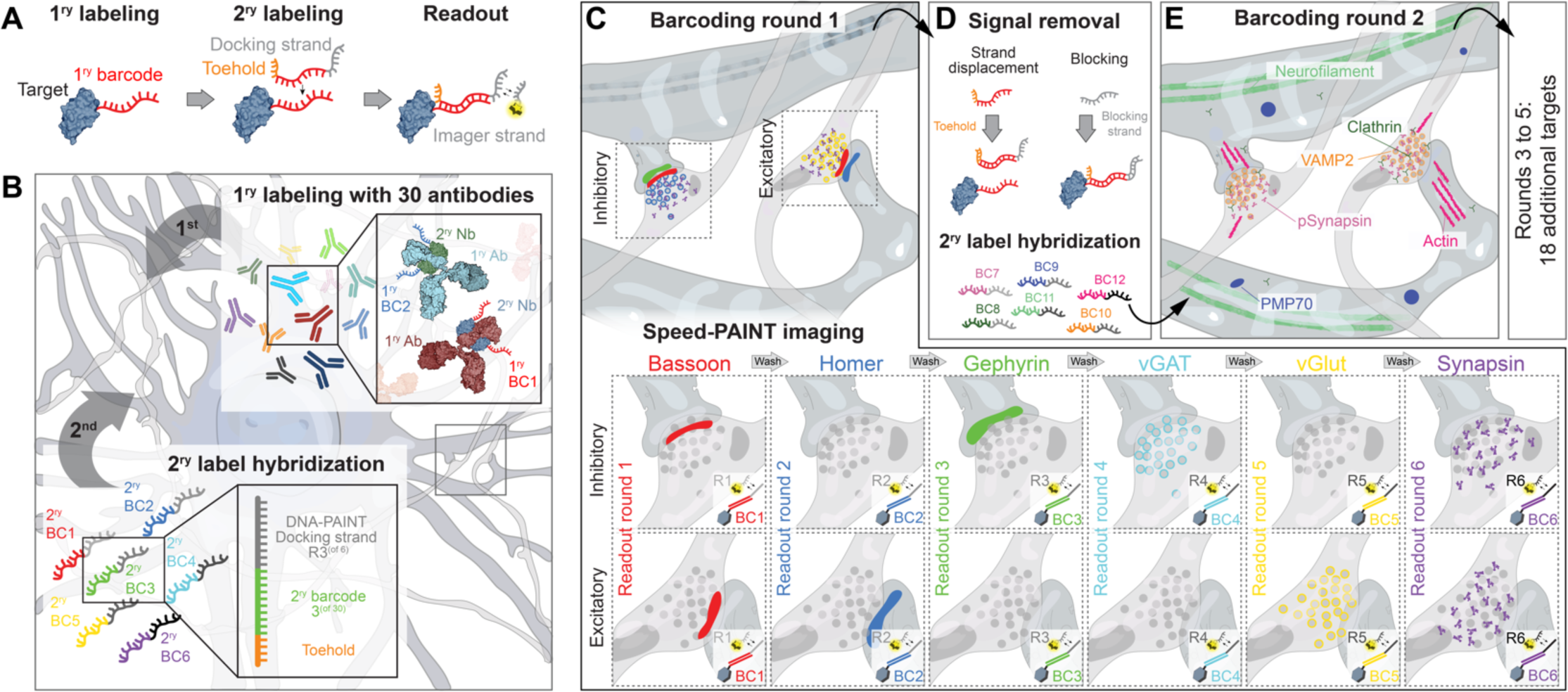
Secondary-label-based DNA-PAINT enables highly multiplexed imaging of single proteins. **(A)** A target of interest is labeled with a primary (1^ry^) DNA barcode (red), followed by a secondary (2^ry^) label, which is composed of the full barcode complement sequence as well as the toehold (orange) and the speed-optimized DNA-PAINT docking sequences (gray). For the readout, a complementary dye-labeled imager strand is added. **(B)** Protein labeling is performed by primary antibodies (Abs) preincubated with secondary nanobodies (Nbs) carrying the 1^ry^ barcodes. Subsequently, a subset of six 2^ry^ barcodes (BC1–BC6) carrying individual Speed-PAINT docking sequences (R1–R6) are hybridized to the respective complementary 1^ry^ barcodes. **(C)** Barcoding round 1 consists of six-target Exchange-PAINT imaging using speed-optimized sequences. In this instance, a subset of both excitatory and inhibitory pre- and post-synaptic proteins are targeted. **(D)** Post acquisition, 2^ry^ labels are inactivated by a combination of toehold-mediated strand displacement and docking strand blockage. Afterwards, a new set of 2^ry^ barcodes (BC7–BC12) again carrying Speed-PAINT sequences (R1–R6) are introduced. **(E)** Barcoding round 2 is then performed similarly to **(C)** and the whole procedure is repeated until all protein targets are acquired (30 total targets in this schematic).

In our implementation of SUM-PAINT, sample preparation starts with the primary labeling step. Antibodies for each cellular target are preincubated with a secondary nanobody carrying a unique primary barcode. Secondary nanobody preincubation allows species-independent immunofluorescence^18^. Labeling of protein targets with these probes is then followed by a secondary hybridization step, where initially six secondary labels (each carrying one out of six available speed-optimized sequences) are hybridized to the respective primary barcodes (Figure 1B). Next, the six targets are sequentially read out using speed-optimized imager strands, as schematically shown for a target region containing an inhibitory and excitatory synapse (R1–R6, Figure 1C). After the first barcoding round is read out, the six secondary labels are removed (Figure 1D) for signal extinction. SUM-PAINT offers two possibilities for this extinction, which can be combined to achieve optimal performance depending on the target molecules: either the secondary label is removed via toehold-mediated strand displacement^17^ or it is blocked via hybridization of a stable 19-nt complement to the DNA-PAINT docking sites. Following signal extinction, the next six secondary labels are hybridized to their respective primary targets, and the process is repeated until all 30 targets have been imaged (Figure 1E).

The SUM-PAINT technique can be adapted for other affinity reagents such as nanobodies or targets like nucleic acids. We note that signal removal by toehold-mediated strand displacement or blocking is a gentler approach to signal extinction compared to most other sequential multiplexing techniques, which depend on the use of relatively harsh denaturation reagents^19^.

## RESULTS

### SUM-PAINT enables speed-optimized multiplexed imaging at sub-5-nm resolution

To effectively apply SUM-PAINT in highly multiplexed, proteome-scale imaging experiments, it is crucial to benchmark its performance concerning spatial resolution, binding kinetics, and signal extinction efficiency. We first employed the programmability of DNA origami nanostructures to quantitatively assess SUM-PAINT’s performance. To achieve this, we designed and assembled two DNA origami structures, each containing 12 binding sites arranged in a 20 nm x 15 nm grid (Figure 2A red sites, Supplementary Table 1). The first nanostructure carried direct DNA-PAINT docking strands, while the second one featured primary barcodes to which secondary labels were subsequently hybridized. Both structures were then simultaneously imaged using the same DNA-PAINT imager strand, followed by a second Exchange-PAINT imaging round for structure identification, targeting a specific geometrical barcode (Figure 2A, yellow sites). Figure 2B represents an exemplary region with two zoom-ins, displaying one direct and one secondary label-extended structure. In both cases, single sites are clearly resolved with high fidelity. A quantitative comparison of the binding kinetics (n = 400 structures) demonstrates that the secondary-label-based DNA-PAINT structures exhibit similar bright times (τb), but intriguingly show a 30 % shorter dark time (τ_d_), leading to an increased image acquisition speed with the same target sampling (Figure 2C). Subsequently, we tested the attainable spatial resolution of secondary-label-based DNA-PAINT and successfully resolved a 5 nm spaced MPI logo structure with a localization precision (σ) of approximately 1.8 nm (Figure 2D), representing current cutting-edge DNA-PAINT spatial resolution^16, 20^. In summary, secondary-label-based DNA-PAINT is on par with (and, in terms of acquisition speed, even superior to) the current state-of-the-art in speed-optimized DNA-PAINT.

**Figure 2.**
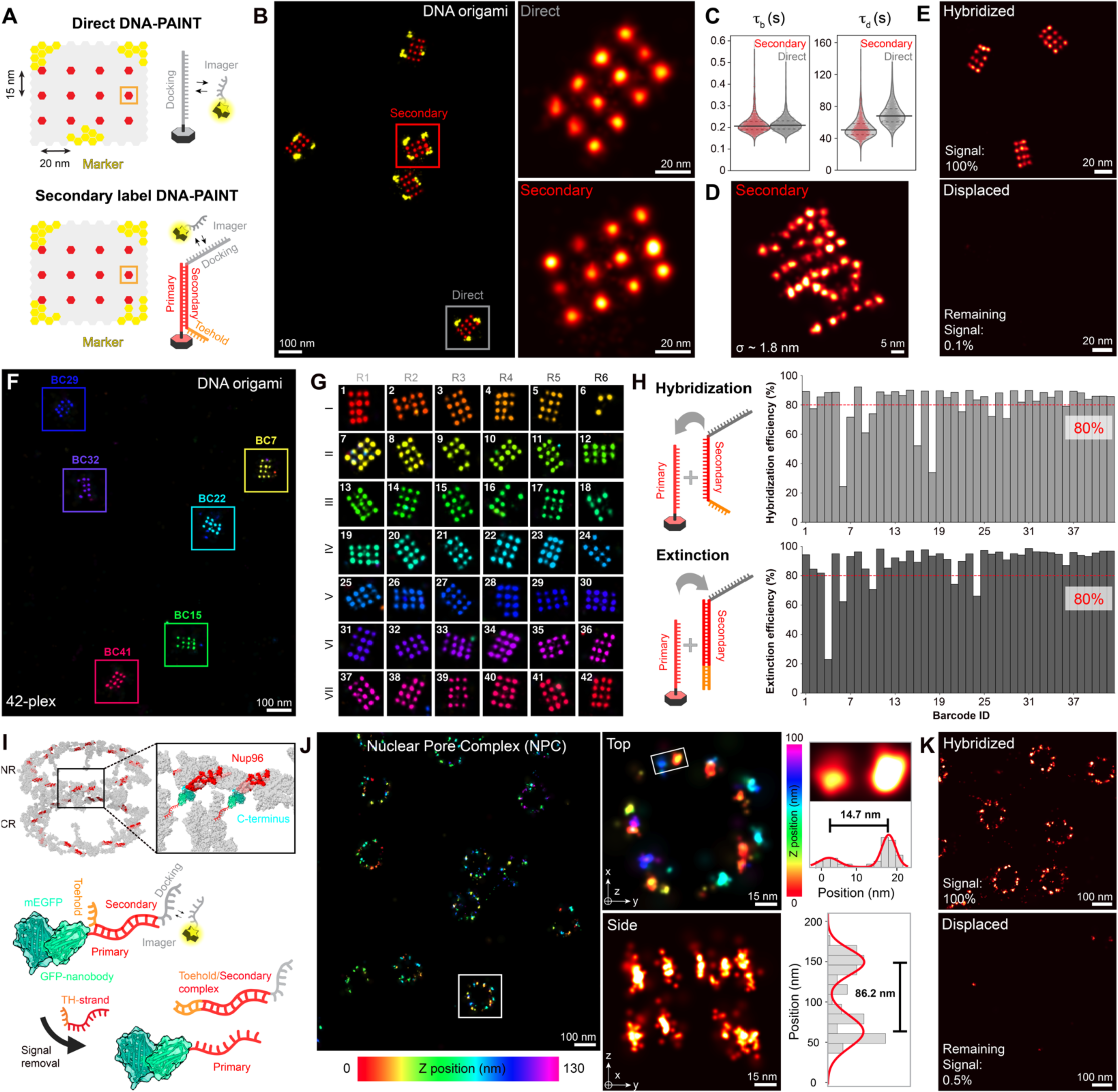
SUM-PAINT enables speed-optimized multiplexed imaging at sub-5-nm resolution. **(A)** Schematic representation of DNA origami structures used for benchmarking direct vs. secondary-label DNA-PAINT. **(B)** Simultaneous acquisition of direct vs. secondary-label DNA-PAINT yields equivalent imaging performance (geometrical barcode (yellow) is acquired using Exchange-PAINT for structure identification). **(C)** Binding kinetic comparison between direct and secondary-label DNA-PAINT shows similar bright times (τb), yet 30% shorter dark times (τd), yielding faster image acquisition. **(D)** Although the binding site is spaced further from the target in the secondary label case, state-of-the-art sub-5-nm image resolution is demonstrated by resolving the 5-nm MPI logo on DNA origami. **(E)** Using toehold-mediated strand-displacement, less than 0.01 % of the signal remains after displacement. **(F)** Representative region of a 42-plex screening experiment showing six different 15-nm DNA origami grid structures, each carrying one out of 42 primary barcodes. **(G)** Codebook for the 42-plex screening experiment with an exemplary DNA origami for each barcode, with columns representing barcoding rounds and rows imager specific readout rounds. **(H)** Hybridization efficiency (a combination of primary barcode incorporation and secondary label hybridization efficiency) for all 42 barcodes (**top**). Displacement efficiency (percentage of correctly displaced secondary labels) for all 42 barcodes (**bottom**). **(I)** The Nuclear Pore Complex (NPC, PDB: 7PEQ) is used as a cellular benchmark for secondary-label DNA-PAINT. Nup96-GFP is labeled with DNA-conjugated anti-GFP nanobodies. **(J)** Exemplary secondary-label 3D-DNA-PAINT overview image color-coded for height shows well-resolved NPC structures. Zoom-in of the highlighted NPC in the overview (**top right**) with cross-sectional histogram fit revealing well-resolved single Nup96 proteins at 14.7 nm distance. Side-view of the highlighted NPC in the overview with cross-sectional histogram fit reveals well-resolved nuclear and cytoplasmic parts of the NPC (**bottom right**). **(K)** Using toehold-mediated strand-displacement, 0.05 % of the signal remains after displacement.

To evaluate the strand removal efficiency for signal extinction, we performed toehold-mediated strand displacement of the secondary label strand on DNA origami. Using an optimized hybridization buffer with 100 nM toehold probes, we successfully achieved a displacement efficiency of over 98 % in less than 2 min (Figure 2E and Supplementary Figure 1A and B).

Next, we investigated SUM-PAINT’s performance in sequential multiplexing. For this purpose, we designed 42 orthogonal DNA origami grid structures, each equipped with a distinct primary barcode sequence (Supplementary Table 3). Figure 2F shows an exemplary field of view of six DNA origami structures with unique barcodes. Figure 2G presents a gallery with a representative structure for each barcode.

From this 42-plex SUM-PAINT experiment, we assessed the hybridization and signal extinction efficiency of secondary labels as two main performance criteria. The quantitative "completeness" of each DNA origami structure served as an indicator for the hybridization efficiency of the secondary label. We obtained an average incorporation efficiency of ∼80 % (with 81 % displaying over 85 %). Since the average incorporation efficiency of strands in DNA origami structures evaluated with direct DNA-PAINT^16^ has been determined to be ∼85 %, SUM-PAINT’s performance is once again on par with cutting-edge direct DNA-PAINT (Figure 2H).

Next, we evaluated the ability to extinguish the signal of the secondary label using toehold-mediated strand displacement. Most barcodes showed displacement efficiency greater than 80 % (with 64 % displaying over 85 %) (Figure 2H). We then used a combined performance metric, considering both hybridization and extinction efficiency, to select the best-performing barcodes for downstream experiments. As a final set of evaluation experiments, we performed cellular benchmarking using the nuclear pore complex (NPC) as a target^21^. The NPC has recently been recognized as a major benchmark structure for super-resolution microscopy^21, 22^. We focused on Nup96, a structural NPC protein that is part of the so-called Y-Complex. Nup96 is present in eight pairs, exhibiting an eight-fold symmetry on both cytoplasmic and nuclear rings, totaling 32 copies per NPC. A homozygous Nup96-GFP knock-in cell line was used along with anti-GFP nanobodies conjugated to a primary sequence for labeling (Figure 2I). We then performed 3D imaging and successfully resolved Nup96 proteins both laterally and axially at the expected distances (Figure 2J).

Additionally, we evaluated the displacement efficiency of secondary labels using toehold-mediated strand displacement in cells. Applying the DNA origami-optimized displacement conditions, we found that a slightly longer displacement time was necessary for efficient strand removal, likely due to the more complex cellular sample environment. However, we were able to achieve a displacement efficiency of 99.5 % after 15 min (Figure 2K).

Taken together, SUM-PAINT facilitates state-of-the-art DNA-PAINT spatial resolution and imaging efficiency i*n vitro* using DNA origami structures and *in situ* with NPCs as test structures. The high performance of SUM-PAINT now enables us to apply it for obtaining a highly multiplexed neuron cell atlas.

### 30-plex neuron atlas at single-protein resolution

Building a neuronal cell atlas with SUM-PAINT necessitates not only the adaptation of cutting-edge imaging from single to multiple targets, but also demands the evaluation of suitable affinity reagents to ensure high specificity and labeling efficiency. Furthermore, multiplexing requires all binders to perform optimally under the same sample fixation and preparation conditions, adding another layer of complexity. Finally, SUM-PAINT multiplexing requires seamless transition between imaging targets without introducing staining or imaging artifacts.

We initially evaluated the performance of more than 100 binders using confocal and STED microscopy, followed by single-target DNA-PAINT. Pre-selected suitable binders were then evaluated for SUM-PAINT multiplexing, resulting in well-performing antibodies for 30 target proteins (Supplementary Table 2). To increase the signal extinction efficiency in SUM-PAINT and thus the overall performance in neurons, we combined toehold-mediated strand displacement with the hybridization of a stable complement to the docking sequence, reducing the remaining post-extinction signal to less than 2 % (Supplementary Figure 1C).

In our current SUM-PAINT implementation, one individual imaging round (comprising one target) in neurons takes approximately 17 min and yields an average localization precision of 6.6 nm, providing label-size-limited single-protein resolution. Adding time for efficient hybridization and signal extinction, a 12-plex SUM-PAINT experiment can be completed in less than 5 hours (Supplementary Figure 2). This represents a significant efficiency gain compared to traditional, non-speed-optimized Exchange-PAINT, which would require more than 300 hours to achieve a comparable spatial resolution and target sampling. While in theory, SUM-PAINT can be scaled to all 30 binders, the challenging neuronal environment requires a more complex staining procedure for higher multiplexing (see Methods). Consequently, the total time required to construct a 30-plex neuronal atlas at single-protein resolution extends to approximately 30 hours. This starkly contrasts with traditionally non-optimized DNA-PAINT, which would necessitate over 800 hours of acquisition, equivalent to more than a month of continuous data gathering, making such an endeavor practically unfeasible.

Figure 3A shows the results of a 30-plex SUM-PAINT experiment, allowing us to map the 3D protein distribution of 30 targets in a single neuron for the first time. Choosing two subsets of protein targets primarily localizing to the cytoskeleton or synapses (Figure 3A, right side) highlights the specificity of SUM-PAINT in this challenging imaging environment. Most excitingly, SUM-PAINT shows specific, highest-resolution signals for each of the 30 proteins when assessed separately (Figure 3B and Supplementary Figure 3).

**Figure 3.**
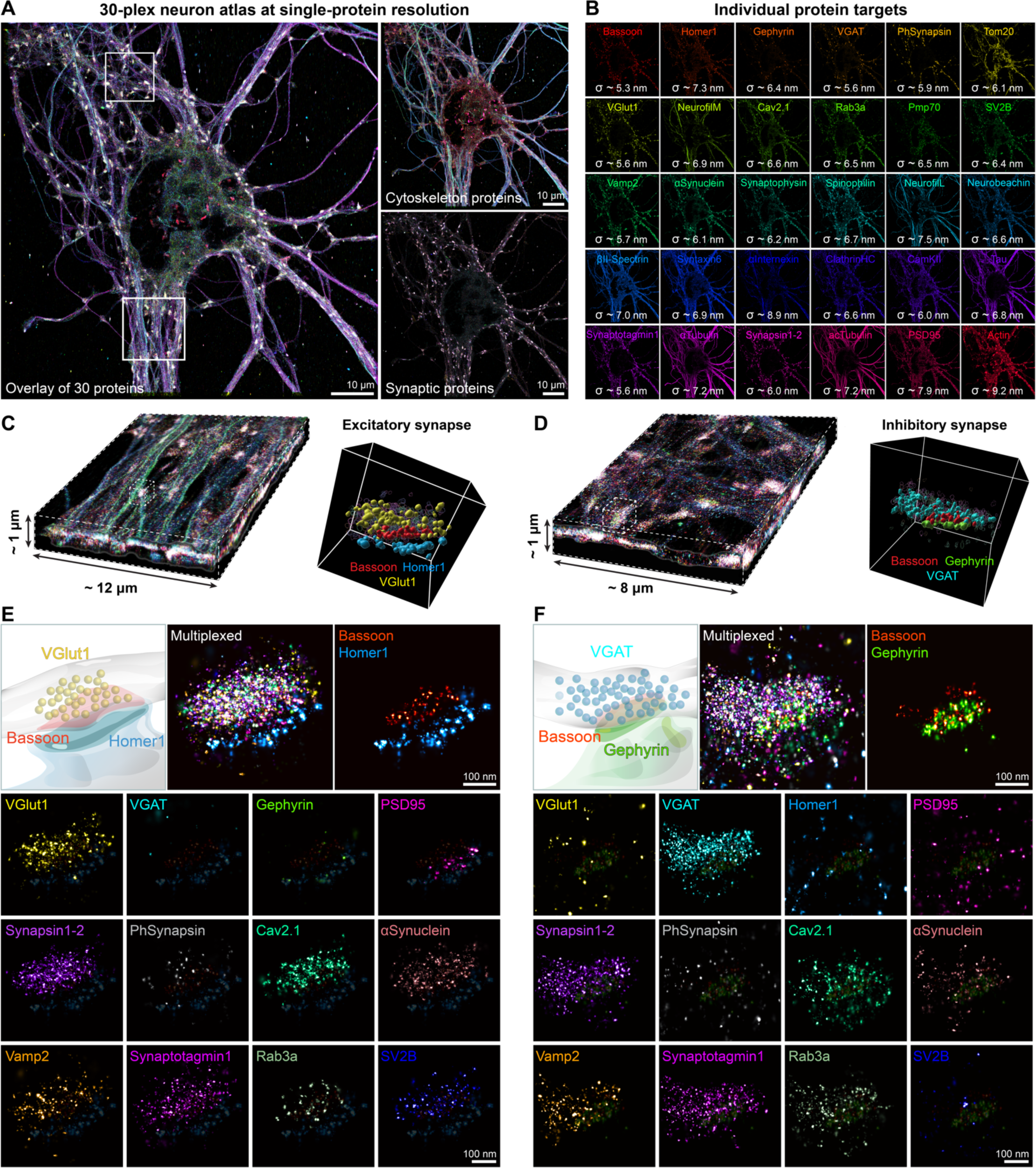
30-plex neuron atlas at single-protein resolution. **(A)** Multiplexed SUM-PAINT overlay image of 30 protein targets visualized simultaneously in an individual neuron with single-protein resolution. Subsets of 9 cytoskeletal (**top right**) and 13 synaptic proteins (**bottom right**) from the same acquisition highlight specificity. **(B)** 30 individual protein targets visualized separately with an average localization precision of 6.6 nm. **(C)** 3D slice of one highlighted region from **A** (**bottom**) alongside a zoom-in of an individual excitatory synapse rotated *en-phase* with the three key proteins Bassoon (red), Homer1 (azure), and VGlut1 (yellow) highlighted. **(D)** 3D slice of the second highlighted region from **A** (**top**) alongside a zoom-in of an individual inhibitory synapse rotated *en-phase* with the three key proteins Bassoon (red), Gephyrin (green), and VGAT (cyan) highlighted. **(E)** The exemplary excitatory synapse from **C** is visualized with all synaptic protein targets in a merged multiplexed view, followed by the scaffold proteins Homer1 and Bassoon as well as all protein targets separately. Note that for visualization purposes, the Bassoon and Homer1 signals are shown at low intensity serving as reference structures for each individual target. **(F)** The exemplary inhibitory synapse from **D** is visualized with all synaptic protein targets in a multiplexed view, followed by the scaffold proteins Gephyrin and Bassoon as well as all protein targets separately. Also in this case, Gephyrin and Bassoon are all shown at low intensity in all overlays.

As we acquire 3D super-resolution data, we can analyze 3D protein distributions in synapses down to the level of individual proteins. Figure 3C displays a 12 x 12 x 1 µm³ imaging volume (region highlighted at the bottom in Figure 3A) along with a zoomed-in view of an excitatory synapse rotated *en-phase*, revealing the composition of the presynaptic scaffold protein Bassoon (red), the excitatory postsynaptic scaffold protein Homer1 (azure), and the excitatory vesicular neurotransmitter transporter VGlut1 (yellow). Figure 3D presents an 8x8x1 µm³ imaging volume (region highlighted at the top in Figure 3A) with a zoomed-in view of an inhibitory synapse, illustrating the nanoscale arrangement of Bassoon (red), the inhibitory postsynaptic scaffold Gephyrin (green), and the inhibitory vesicular neurotransmitter transporter VGAT (cyan). A comparison of the two selected synapses highlights that specific markers for each synapse type, such as Homer1, VGlut1, and PSD95 for excitatory synapses, and Gephyrin and VGAT for inhibitory synapses, display signals exclusively in their respective types (Figure 3E and F). This emphasizes the specificity and efficiency of SUM-PAINT in this complex neuroscientific application. Common synaptic vesicle pool proteins are equally expressed in both types, as is the presynaptic calcium channel Cav2.1, while both synapse types exhibit minimal to no Synapsin I phosphorylation at Ser 549 and only the excitatory type expresses SV2B proteins.

### Multiplexed SUM-PAINT provides nanoscale insights into neuronal architecture

Taking advantage of SUM-PAINT’s exceptional multiplexing and spatial resolution abilities in neurons, we investigated the nano-architecture of various protein species within our highly multiplexed datasets. Initially, for benchmarking purposes, we concentrated on previously-studied structures, analyzing the organization of the cytoskeletal protein Spectrin (βII-Spectrin) and the mitochondrial outer membrane marker Tom20 (Figure 4A). Figure 4B displays a magnified view of the periodic ring-like arrangement of Spectrin, along with an intensity projection. Subsequent Fourier transform and autocorrelation analysis revealed a distance of 190 nm between the rings, consistent with previous studies^24^. A closer look at the distribution of Tom20 localizations revealed the 3D architecture of the mitochondrial outer membrane with individual proteins clearly resolved (Figure 4C). Next, we examined the characteristics of Clathrin-coated vesicles and pits that contribute to the endocytic activities of neurons. Figure 4D shows a representative region focusing on Clathrin heavy chain (ClathrinHC) alongside acetylated tubulin (ac-Tubulin). Closer examination of a single vesicle unveils the circular arrangement of individual proteins forming a spheroid. By aggregating the localizations of over 200 individual vesicles and fitting the intensity maxima of the cross-section of the rings with a two-component Gaussian function, we determined an average diameter of ∼60 nm for the Clathrin-coated pits observed in this region.

**Figure 4.**
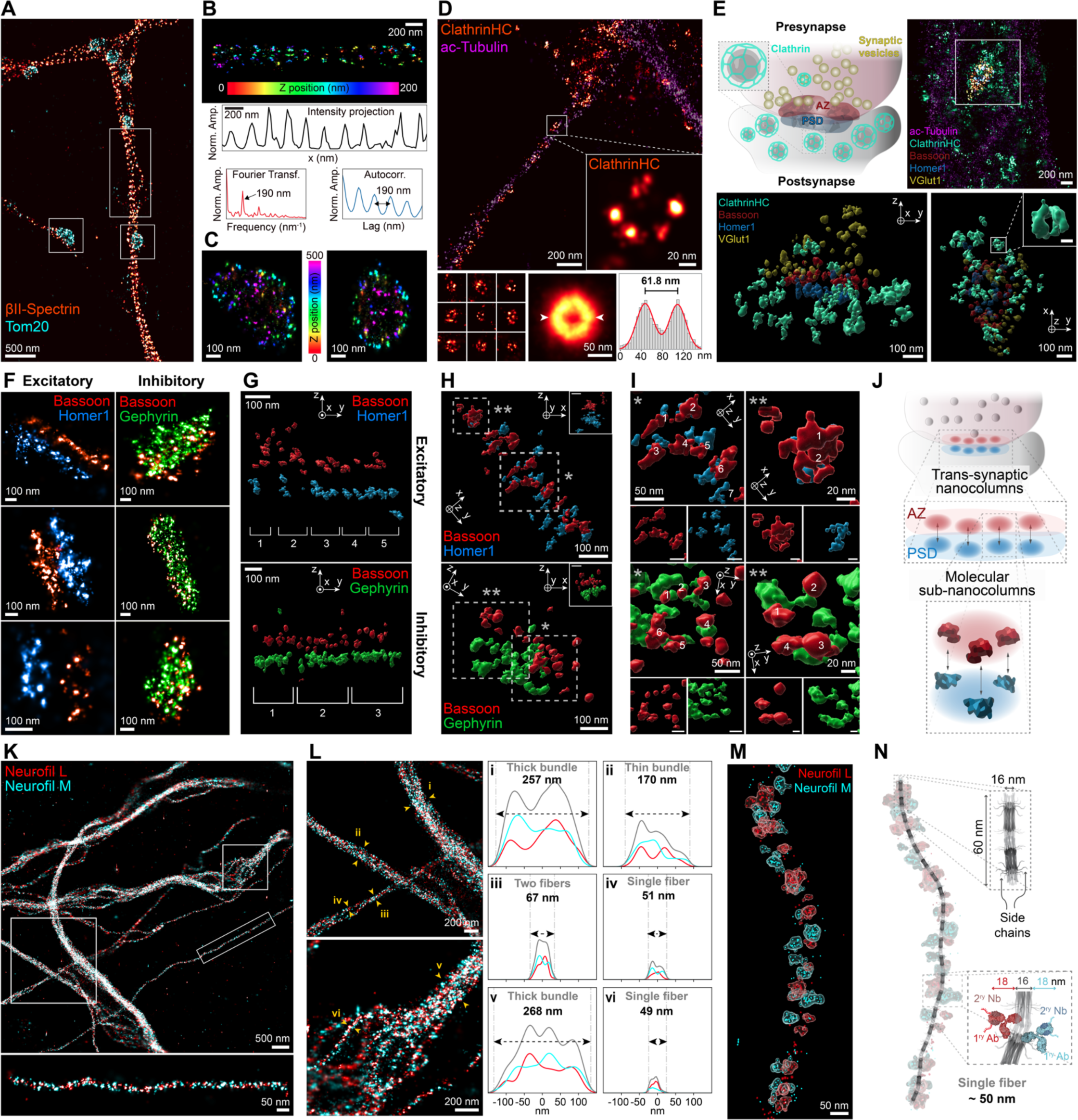
Multiplexed SUM-PAINT reveals nanoscale insights into neuronal architecture. **(A)** 2-plex overlay reveals periodic arrangement of βII-Spectrin (hot look-up table) alongside mitochondria (Tom20, cyan). **(B)** Zoom into the selected spectrin region from (A), color-coded for height. Cross-sectional histogram alongside autocorrelation and Fourier analysis confirms the periodic arrangement of well-resolved Spectrin rings at 190 nm spacing. **(C)** Zoom into highlighted mitochondria from **(A)** reveals single resolved Tom20 molecules (color indicates height). **(D)** 2-plex overview of clathrin alongside acetylated Tubulin as cytoskeleton marker. Zoom-in shows a single traveling vesicle with individual clathrin proteins resolved. Summing of single clathrin vesicles and cross-sectional analysis of the sum image reveals an average diameter of ∼60 nm. **(E)** Top left illustrates a synapse with prominent endocytosis occurring as shown by clathrin vesicles (mint) present in both pre- and post-synapse alongside the Active Zone (AZ) in red, postsynaptic density (PSD) in blue, and synaptic vesicles in yellow. Top right shows a 5-plex image of a representative excitatory synapse. Bottom shows the synapse in volumetric surface rendering in two rotated views, revealing spherical clathrin vesicles (Scale bar: 50 nm). **(F)** Exemplary excitatory and inhibitory synapses showing spatial correlation between pre- and postsynaptic scaffolds. **(G)** Excitatory and inhibitory scaffolds rotated *en-phase* and shown as volumetric surface rendering reveal the molecular arrangement of pre- and postsynaptic scaffolds into nano-pillars. **(H, I)** Upon closer inspection, pre- and postsynaptic overlays reveal that in fact individual nano-pillars exhibit molecular matches for single proteins (numbers indicate individual matches). **(J)** Illustration of trans-synaptic nanodomains (AZ, red; PSD, blue). Not only is the scaffold arranged in small nanopillars (top), but inside the domains individual protein pairs form a molecular match (bottom). **(K)** 2-plex overlay of intermediate neuronal filaments (medium **(M)** and light **(L)** chain in cyan and red respectively). Zoom-in shows a selected individual filament with alternating medium and light chains (bottom). **(L)** Two exemplary selections indicating the arrangement of neurofilament into different strength bundles. Top shows a selection of different thicknesses, while bottom shows the organization of 8 individual filaments into a thick bundle. Line profiles of respective filaments (i-vi), with respective thicknesses conserved across the sample. **(M)** Zoom-in of an individual fiber rendered as localizations and volumetric surface, showing individual protein molecules. **(N)** Illustration of the proposed neurofilament structure from electron microscopy (EM)^23^ labeled with antibodies. Two antibodies with an average size of 18 nm in addition to the 16 nm filament size result in an approx. 50 nm fiber. Remarkably, the single and two fibers from **(L)** are consistent with the thickness measured by EM.

In addition to examining the distribution of individual clathrin pits along the dendrites and axons of neurons, we were interested in the endocytic events that occur at synapses. Figure 4E shows clathrin alongside the active zone, the postsynaptic density, and synaptic vesicles with an exemplary image of an excitatory synapse with prominent endocytosis occurring in a 5-plex overlay. An *en-phase* arrangement of volumetric surface-rendered localizations revealed two clathrin vesicles in the presynapse and most vesicles in the postsynapse, as indicated by the position of the scaffold proteins Bassoon and Homer1 and the excitatory vesicle pool marker VGlut1. By rotating the synapse, we extracted a single clathrin vesicle from the postsynapse, measuring ∼100 nm in size. To further explore the neuronal nanoarchitecture, we examined the previously-observed assembly^25^ of pre- and postsynaptic scaffolds into trans-synaptic nanocolumns (Figure 4F). We clearly observed juxtaposed localization clusters ranging from 50 to 100 nm in the pre- and postsynaptic scaffolds, illustrating the nanocolumnar substructure of the adjacent scaffolds (Figure 4G), as previously reported^25^. Since DNA-PAINT provides single-protein resolution, we took a closer look at these structures by rotating the scaffold to observe the alignment of individual nanocolumns. Figure 4H displays the previously selected excitatory synapse alongside an additional inhibitory synapse. Further zooming into the nanocolumns and separating the pre- and postsynaptic protein distributions revealed that the scaffold is in fact aligned at the level of individual pre- and postsynaptic proteins for both synapse types and across multiple regions (Figure 4I). Interestingly, the macromolecular assembly across synapses appears not only to occur in the columnar arrangements, but these structures seem to be correlated down to the level of single proteins, potentially forming a stoichiometric molecular match that could indicate transsynaptic macromolecular complexes (Figure 4J).

The final structure we examined was the arrangement of neuron-specific intermediate filaments, the neurofilaments, with a particular focus on the light and medium chains of these polymers. Figure 4K shows an exemplary 2-plex composition of the light and medium chains of the neurofilament, with a zoom-in on a single fiber at single-protein resolution, revealing alternating polymers. We further demonstrated that individual neurofilament fibers are organized into bundles of varying thickness and assembled into higher-order structures containing up to eight individual fibers (Figure 4L). A cross-sectional profile of selected regions revealed that both the individual fiber width of approximately 50 nm and the thickest bundle width of approximately 260 nm appeared to be well conserved throughout the sample. Next, we explored the molecular arrangement of these neurofilaments. Figure 4M shows a close-up of an individual fiber. We determined a fiber thickness of approximately 50 nm from our DNA-PAINT images, which aligns well with EM-derived measurements^23^ of 16 nm, given the overall size of our labeling probe complex of approximately 18 nm (Figure 4N). This is further supported by the fact that the thickness of two fibers increased by 16 nm compared to a single fiber.

While DNA-PAINT’s single-protein resolution offers unparalleled capabilities for investigating the nanoarchitecture of neurons, its extensive multiplexing provides large and complex multidimensional datasets that require sophisticated analysis tools for full exploitation. The power of SUM-PAINT, with its multiplexing and 3D super-resolution capabilities, is that it allows us to map the structure and shape of protein assemblies, place them in context with other proteins, and access a vast parameter space for exploration. This is ideal for studying the heterogeneity of individual synapses.

### Machine learning applied to spatial proteomics at single-protein resolution reveals a new synapse type

Next, we shifted our focus to the composition of synapses, as these are central structures for proper brain function and their complexity has not been explored at this level of resolution and multiplexing. With an atlas of 30 protein species in six independent datasets (Supplementary Figure 4) at our disposal, our aim was to uncover the intricate details of synaptic protein compositions and diversity. However, due to the spatial and highly multiplexed complexity of the data, this is a challenging task.

UMAPs are frequently used for analyzing highly multiplexed transcriptomic, genomic datasets, and diffraction-limited images of protein distributions^26^. However, our SUM-PAINT approach takes this a step further by using single-protein resolution super-resolution imaging data, which enriches the available parameter space by adding morphometric features to protein identity. SUM-PAINT enables 3D nanoscale mapping of biomolecules, assessing characteristics like shape, density distribution, or protein count, while spatial cluster analysis further delineates surfaces, volumes, and correlations between protein species.

To effectively use the vast features space that SUM-PAINT offers, we developed an integrated workflow for examining highly multiplexed super-resolution data at single-protein resolution. Figure 5A outlines our approach to data extraction and analysis. Initially, individual synapses were selected based on pre- and postsynaptic protein signals (see Methods for details). Subsequently, 1590 unique features from each synapse were extracted by generating 3D histograms of the protein channels, followed by a DBSCAN spatial cluster analysis. A comprehensive list of the total number of extracted features for each synapse, as well as their respective values, is provided in Supplementary Table 4. Ultimately, all features were consolidated into a UMAP, and feature-space clustering was applied (Figure 5B).

**Figure 5.**
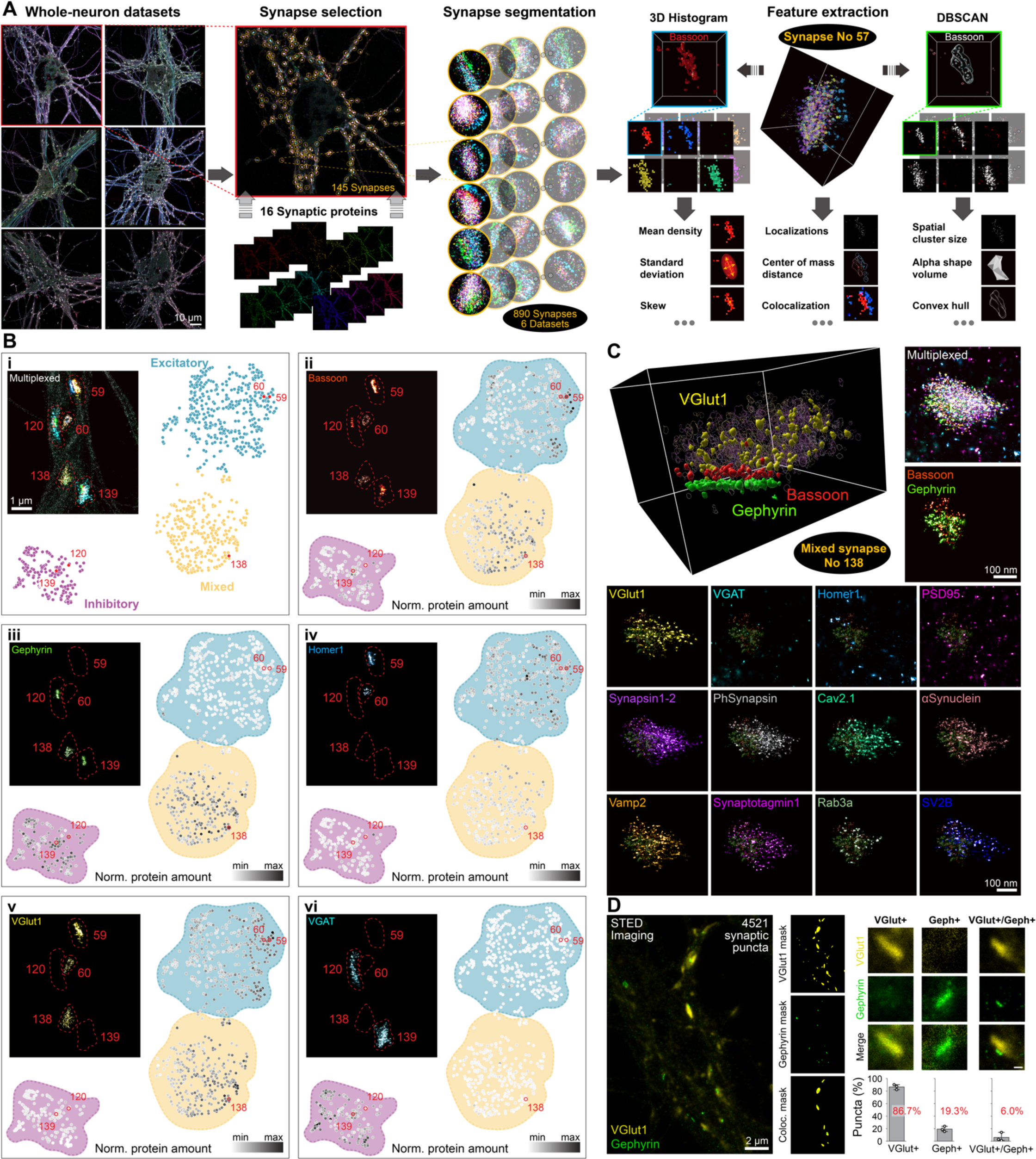
Machine learning applied to spatial proteomics at single-protein resolution reveals a new synapse type. (**A**) Feature extraction workflow of six multiplexed whole-neuron datasets. In total 890 synapses are segmented from a 16-plex overlay of synaptic proteins. The localizations in the segmented synapses are analyzed as 3D histograms and DBSCAN spatial clustering for parameter extraction, resulting in 1590 features per synapse. **(B)** A UMAP of 1590 features is created and clustered (**i**), yielding three populations of synapses. The individual synapses are labeled with their identification number from the analysis workflow and are highlighted in the UMAP. The top left shows the multiplexed overlay of excitatory and inhibitory markers alongside acTubulin (dark green background) as cytoskeleton reference. The unique composition of pre- and postsynaptic markers (Bassoon, Homer1, Gephyrin), as well as the neurotransmitter transporter (VGlut1, VGAT) allows us to label the UMAP clusters as excitatory (blue), inhibitory (purple) and a new mixed synaptic population (yellow). (**ii–vi**) highlights the normalized protein content (grayscale) of a single pre- or postsynaptic marker in the UMAP. The top left shows a single channel image of the same region for each protein. **(C)** Volumetric surface rendering of a mixed synapse rotated *en-phase* (top left), containing VGlut1 as a neurotransmitter transporter, Bassoon as presynaptic, and Gephyrin as postsynaptic marker. Top right represents the same synapse in a 13-plex overlay and as a 2-plex image containing Bassoon and Gephyrin. The bottom shows 12 additional protein channels at single-protein resolution. **(D)** High-throughput STED imaging of more than 4520 synaptic puncta indicates that the relative abundance of the mixed synapse type is 6 %.

Interestingly, we identified three major clusters, going beyond the conventional distinction between excitatory and inhibitory synapses in these cells. To investigate this "unexpected" synapse type, we color-coded the UMAP based on the expression level of key proteins that determine inhibitory and excitatory synapses: Bassoon as a general presynaptic scaffold, Gephyrin as an inhibitory postsynaptic scaffold, Homer1 as an excitatory postsynaptic scaffold, VGlut1 as a transporter for the excitatory neurotransmitter glutamate, and VGAT as a transporter for the inhibitory neurotransmitter (γ-aminobutyric acid; GABA).

For further clarification, we traced back synapses from the UMAP analysis and visually examined protein composition for some exemplary synapses (Figure 5B, red) in all channels corresponding to different protein types. As an example, by visualizing the protein content, we concluded that synapses expressing Homer1 and VGlut1 (e.g., synapse 59 and 60) belong to the excitatory type, while synapses expressing Gephyrin and VGAT (e.g., synapse 120 and 139) belong to the inhibitory type.

Intriguingly, the synapse with identification number 138 belongs to a previously unreported type, which we define as "mixed”. The mixed type is characterized by comprising VGlut1 as a neurotransmitter vesicular transporter, originating from a glutamatergic neuron, but coupled with a postsynapse expressing gephyrin as a scaffold protein.

Next, we visualized synapse 138 using volumetric surface rendering and examined the protein composition of this “mixed” synapse by rendering each protein channel individually (Figure 5C). There was no significant signal for either Homer1 or VGAT, while both Gephyrin and VGlut1 displayed clearly defined spatial protein clusters, aligning with the synapse morphology at the nanomolecular level. Extending these findings to all datasets, based on the SUM-PAINT analysis, we deduced that mixed synapses could be identified by the concomitant positive signals in the VGlut1 and Gephyrin channels, but by the absence of Homer1 (Supplementary Figure 5A for more examples). We also studied the overall abundance of the mixed synapses by two-color STED microscopy, which allowed us to characterize over 4,500 synaptic puncta (Figure 5D). Analysis of these puncta revealed that approximately 6 % were of the mixed type. To corroborate our findings in a broader context we uncovered the mixed synapses in brain slices using STED imaging (Supplementary Figure 5B).

To further elucidate the nature of these synapses, we conducted an assessment of potential developmental influences on synapse numbers within our culture model (refer to Supplementary Figure 5C), as well as the distribution of different neurotransmitter receptors (see Supplementary Figure 5D). These experiments indicated that there is no significant developmental influence on the percentage of this type of synapses, at least in these cultures and that the postsynaptic receptors are, in 80 % of the cases, of the Aγ-2 GABAergic type.

The discovery of the mixed synapse type exemplifies the unique capabilities of combining highly multiplexed super-resolution microscopy with machine learning-based analysis for data exploration. While the significant differences in protein composition between excitatory, inhibitory, and mixed synapses led to three pronounced clusters in the UMAP, the extensive feature space could offer further insights into synapse subtypes and their statistical and morphometric features, which we subsequently examined in more detail.

### High-content feature space analysis enables deep investigation of synapse type characteristics revealing synaptic diversity

To delve deeper into synaptic features, we extended our analysis to include all synaptic proteins (Figure 6A). In addition to the pre-(Bassoon) and postsynaptic scaffold (PSS: Homer1, Gephyrin) proteins and neurotransmitter transporter proteins (NTT: VGlut1, VGAT), the analysis also incorporates several synaptic vesicle and vesicle pool proteins (Synaptotagmin1, Rab3a, Vamp2, SV2B), vesicle pool associated proteins (Synapsin, Phospho-Synapsin, α-Synuclein), and the calcium channel protein Cav2.1 (Figure 6B). To further understand the characteristics of different synapses and their heterogeneity, we focused on the following features: i) Center of Mass (CoM) position for each protein localization cluster, ii) Spatial cluster volume for each protein species, iii) Distance profile within the synapse types, iv) Volume correlation and hierarchical clustering between protein species.

**Figure 6.**
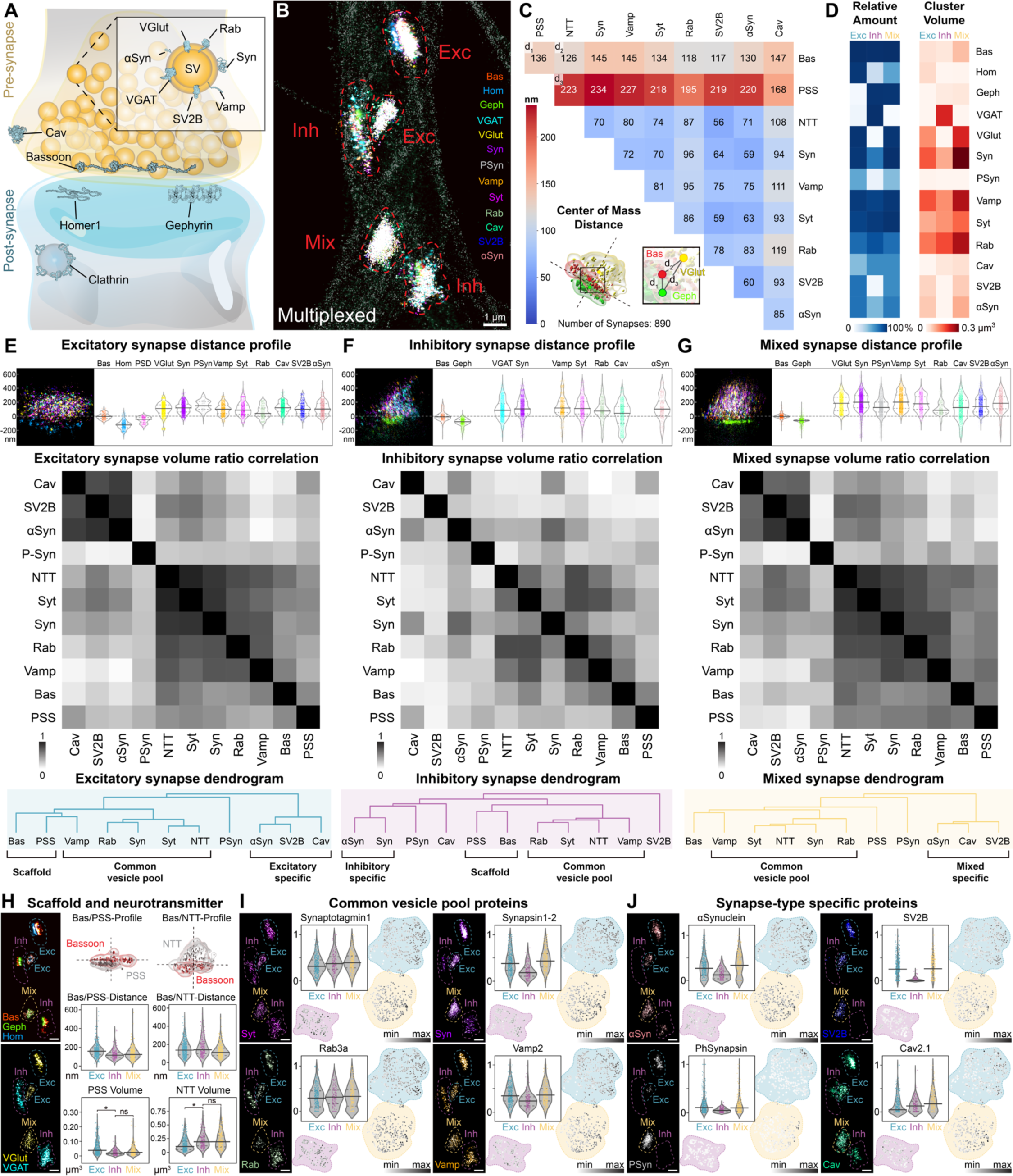
High-content feature space analysis enables deep characterization of synaptic diversity. **(A)** Schematic illustration of synaptic proteins included in the high-content analysis. Note that for representation purposes VGlut1 and VGAT as well as Homer1 and Gephyrin are shown in the same scheme. **(B)** Multiplexed super-resolution overlay of 13 synaptic proteins under investigation. **(C)** Average center-of-mass (CoM) distance matrix of 890 synapses alongside illustration of CoM analysis of three exemplary synaptic proteins with respective distances d1, d2, and d3 (bottom left). **(D)** Left: Heatmap of the normalized relative protein counts for each protein type for excitatory, inhibitory and mixed synapses. Right: Protein cluster volume (in μm^3^) for each protein type for excitatory, inhibitory and mixed synapses. **(E)** Top: Exemplary excitatory synapse localization distribution after DBSCAN spatial cluster detection. Center: Heatmap of the Pearson correlation coefficients of the protein volume ratios. Bottom: Dendrogram resulting from a hierarchical cluster analysis of the volume ratio correlation of all synaptic proteins. **(F)** Same as **(E)** for inhibitory synapse type. **(G)** Same as **(E)** for mixed synapse type. **(H)** Analysis details for scaffold and neurotransmitter transporter (NTT) proteins. Top left shows a 3-plex composition of scaffold proteins with the synapse type indicated. Bottom left shows the respective region for the NTT. Top middle and right show an exemplary profile of the presynaptic scaffold protein Bassoon (Bas) to the postsynaptic scaffold (PSS) (middle) and NTT (right) with volumetric rendering. Center middle and right show the synapse type specific CoM distance for Bas to the PSS (middle) and to the NTT (right). Bottom middle and right show the synapse type specific protein volume of PSS and NTT. **(I)** Analysis details for the common vesicle pool proteins. Top left shows the individual imaging result for Synaptotagmin1 of the selected region alongside the relative normalized protein content represented as average in the synapse type specific violin plot and as grayscale colored UMAP. Top right, bottom left and right display the same arrangement for Synapsin1-2, Rab3a and Vamp2. **(J)** Analysis details of synapse type specific proteins. Same as **(I)** for α-Synuclein, SV2B, Phospho-Synapsin at serine 549 (site6; PhSynapsin) and Cav2.1. In this figure Protein names were abbreviated the following way: (Bassoon:Bas, Homer1:Hom, VGlut1:VGlut, Synapsin1-2:Syn, PhSynapsin:PSyn, Vamp2:Vamp, Synaptotagmin1:Syt, Rab3a-Rab, Cav2.1:Cav, α-Synuclein:αSyn, Gephyrin:Geph)

The first feature we examined in greater detail was the center of mass distance (CoM) of the spatial protein clusters. Comparing CoM distances between protein species enabled us to extract metrics such as colocalization and average distance. We represented the CoM distance between all proteins (obtained from an average of 890 synapses) in a 9x9 matrix, highlighting the synapse-specific proteins (PSS and NTT) as individual values (Figure 6C). Our analysis yielded, for example, a distance of 136 nm between Bassoon and PSS, which aligns well with reported data for pre- and postsynaptic distances^27, 28^. Furthermore, the CoM analysis of vesicle pool proteins toward each other reveals much closer distances in the range of 50–80 nm. Given the substantial size of the vesicle pool on the order of microns, this practically indicates colocalization of proteins. For an even more detailed analysis, we also calculated the CoM distances for individual types of synapses in Supplementary Figure 6.

Next, we calculated relative protein amounts and cluster volume for each synapse type (Figure 6D). The matrix displaying relative protein amounts replicates the distributions between PSS and NTT observed from the localization number UMAP analysis in Figure 5, with excitatory synapses displaying Homer1 and VGlut1, inhibitory Gephyrin and VGAT, and mixed Gephyrin and VGlut1. Additionally, we can infer that the common vesicle pool proteins (Synaptotagmin1, Rab3a, and Vamp2) exhibit similar expression levels, whereas Synapsin, PhSynapsin, SV2B, and α-Synuclein are present in lower amounts in the inhibitory synapses. In terms of volume, pre- and postsynaptic scaffolds display an average volume of 0.05 µm³, while the vesicle pool proteins form a much larger structure with volumes of approximately 0.3 µm³.

The top panel of Figure 6E-G presents an exemplary image of an individual synapse alongside its protein localization profiles as violin plots, after spatial cluster detection. The profile reveals that pre- and postsynaptic scaffolds for all three synaptic subtypes are spaced approximately 150 nm apart, while PSD95 is localized closer to the presynaptic site than Homer1 for the excitatory synapse.

To evaluate if and which specific protein species are correlated in each of the three synaptic subtypes, we first calculated the relative volume ratio between each protein species for each synapse. After that, we employed a Pearson’s correlation plot to extract the correlation for each protein species. We then compiled the values in an 11 x 11 matrix with the PSS and the NTT specific to the synaptic subtype. The correlation matrices reveal distinct visual differences between excitatory and inhibitory synapses (Figure 6E and F, middle).

To assess this information more quantitatively, we employed hierarchical clustering to construct a dendrogram illustrating the similarity clusters of protein species (Figure 6E and F, bottom). Pre- and postsynaptic scaffold proteins (Bassoon and PSS, respectively) exhibit a strong correlation for both excitatory and inhibitory synapses, as do the common vesicle pool proteins Synaptotagmin 1, Rab3a, Vamp2, and the NTT. Interestingly, in excitatory synapses, Synapsin demonstrates a strong correlation with other common vesicle pool proteins, while for inhibitory synapses, it exhibits a strong correlation with α-Synuclein, forming an inhibitory-specific cluster. Notably, α-Synuclein, SV2B, and Cav2.1 display a pronounced correlation cluster in excitatory synapses, while no correlation is observed in the inhibitory case, making it an excitatory-specific cluster. Phosphorylation of Synapsin at site 6 (Ser 549), which we term PhSynapsin, does not correlate with any other protein, indicating that future more detailed studies of post translational modifications events will broaden our knowledge about synaptic diversity.

We then compared the volume ratio correlation matrix and dendrogram of the mixed to those of excitatory and inhibitory synapses (Figure 6G). The mixed type exhibits greater similarity to excitatory synapses than inhibitory ones. This is probably also due to the fact that the proteins that we analyzed are predominantly presynaptic and the neurotransmitter for the mixed synapse type is glutamate (indicating an excitatory origin).

While the volume correlation analysis between proteins among the three synaptic subtypes highlights the similarity between excitatory and mixed synapses and the difference with inhibitory synapses, our feature space provides a more profound opportunity to investigate more general characteristics of the three populations, such as distances between pre- and postsynaptic components across types. Initially, we focused on differences in PSS and NTT between the synapse types by analyzing the distance and volume between the pre- and postsynaptic scaffold and between the presynaptic scaffold and NTT (Figure 6H). The CoM distance comparison plot for the individual synapse types reveals that the distance between vesicles and either pre- or postsynaptic scaffolds depends on the respective PSS.

The volumetric comparison between the PSS of three subtypes also indicates that excitatory Homer1 PSS differs in size from the conserved inhibitory and mixed Gephyrin PSS size. Although NTTs vary from excitatory to inhibitory synapses, the distance between Bassoon and NTT appears conserved. However, the NTT volume, which approximates the size of the synaptic vesicle pool for each synapse, seems to be correlated with the PSS identity, with the excitatory VGlut1 exhibiting a smaller volume than VGAT or VGlut1 in either inhibitory or mixed synapses.

Lastly, we explored how individual correlation clusters resulting from the dendrogram differ on a single-synapse level. We display the average protein expression level as a violin plot and intensity-colored UMAP next to an exemplary region from the imaging data. Our initial target was the distribution of common vesicle pool proteins Synaptotagmin1, Rab3a, Vamp2, and Synapsin. Figure 6I reveals that Synaptotagmin1 and Rab3a are evenly distributed among all three synapse subtypes. Interestingly, Vamp2 and Synapsin show a difference in inhibitory synapses. While Vamp2 is slightly lower expressed in inhibitory synapses, Synapsin exhibits a much lower expression level in both the average overall synapses and the UMAP, where each synapse is shown individually. The lower expression level of Synapsin has been reported before^29^, and our extraction pipeline could replicate this finding.

Our further analysis focused on the synapse-type-specific protein correlation clusters, which indicated a significant difference between synapse subtypes in the volume correlation matrix. Figure 6J shows that α-Synuclein is strongly downregulated in the inhibitory synapses, consistent with previous reports^30^. SV2B exhibits almost no signal in inhibitory synapses, as expected^31, 32^. Moving on to PhSynapsin and Cav2.1, we observed that phosphorylation of Ser 549 is only occurring at excitatory and mixed synapses. Furthermore, Ser 549 phosphorylation on synapsin in these synapse types is a distinct event, suggesting an orthogonal level of regulation, as also clearly indicated by a strong difference in the UMAP intensity color for individual synapses. Cav2.1 appears to be present in all synapse subtypes, and the average reveals that the expression level is lower in excitatory synapses. However, examining the UMAP suggests that Cav2.1 also displays a distribution into distinct events, with all three synaptic subtypes featuring high variability in its expression level.

## Discussion

With SUM-PAINT, we have pioneered a method to advance state-of-the-art super-resolution imaging, achieving unparalleled spatial resolution and throughput for a virtually unlimited number of protein targets. By developing an analysis workflow that incorporates advanced machine learning and omics-inspired approaches, we successfully mapped the multiparameter features of highly multiplexed, high-resolution datasets and zoomed into individual sample regions to reveal nano-arrangements at the level of single proteins. Applying SUM-PAINT to neurons, we generated the most extensive multiprotein dataset to date, at spatial single-protein resolution, comprising up to 30 distinct protein targets in parallel with an average localization precision of 6.6 nm. These highly multiplexed neuronal architecture maps allowed us to resolve spectrin rings, clathrin pits, and synapses with exceptional spatial resolution and open the field to the possibility of analyzing synaptic and even subsynaptic variability in response to neuronal plasticity modulations. Excitingly, we were able to unveil the nano-arrangement of neurofilaments and demonstrated that individual trans-synaptic nanocolumns exhibit a precise molecular alignment on a single-protein level.

To our knowledge, our work is the most comprehensive spatial study to date of the nanomolecular diversity of synapses combining the information of protein identity and detailed morphometric features. The most important aspect of our analysis is that by considering multiple proteins in parallel along with their morphological features, the available parameter space for analysis is expanded, allowing for a comprehensive analysis of synapse diversity. Through our AI-guided, omics-inspired analysis, we have unveiled three distinct synaptic subtypes within hippocampal neurons: the canonical glutamatergic excitatory and GABAergic inhibitory synapses, alongside a novel mixed synaptic subtype. This intriguing mixed subtype contains an excitatory glutamatergic synaptic vesicle pool (VGlut1^+^) paired to a Gephyrin-positive postsynaptic scaffold, typically associated with inhibitory synapses. This newfound synapse category, albeit a minor subset, would have remained hidden without our robust, high-resolution image acquisition and analysis methodology. Given that Gephyrin, to our knowledge, should not directly cluster glutamatergic receptors – a fact we have verified (Supplementary Figure 5D) – it is probable that these synapses do not take part in traditional neurotransmission.

The discovery of this subtype raises a number of additional questions that will form the basis of future research aimed at understanding the physiological significance of these synapses, their possible transient nature, and their role in development and potential disease. At present, our data, which suggests a higher similarity between mixed and excitatory synapses, allows us to hypothesize that these may be chemically inactive synapses, potentially transitioning to a fully excitatory state upon refinement, possibly through a multitude of steps that are still to be discovered.

Moreover, we were able to characterize the details of excitatory, inhibitory, and the newfound mixed synapse type, leveraging a broad range of features. These encompass interprotein cluster distances, shape and volume of protein distribution, as well as their correlation matrices. This analysis ultimately allowed us to decode the specifics of synaptic subtypes. Certain proteins, including alpha-synuclein, SV2B, and Cav2.1, display varying correlations with different synaptic subtypes, possibly due to differential expression in neuronal types and suggesting a differential role in human pathologies. In our exploratory studies, we employed an affinity reagent targeting phospho-Ser 549 of synapsin (PhSynapsin), a synaptic readout of MAP kinase activity, which illuminates variations in synaptic plasticity tied to local synaptic signaling. As anticipated, PhSynapsin displayed clear presynaptic localization akin to total synapsin, suggesting potential functional state variations across synapses. Future investigations that combine modulation of synaptic activity with time-resolved analysis of multiple post-translational modifications will help elucidate how signaling influences synaptic heterogeneity.

As revealed by UMAP distributions (Figure 6I-J), distinct synaptic components exhibit specific gradients across synaptic subtypes. This pattern may indicate a thus far poorly understood temporal evolution of synaptic identities, or alternatively hint at the existence of additional, yet-to-be-discovered subtypes amenable to targeted genetic or pharmacological interventions. Altogether, our findings underscore the vast nanomolecular diversity of synaptic subtypes, transcending mere differences in molecular composition. Our work not only unveils a new synaptic connection subtype, but also implies a unique, uncharted diversity in the action mechanisms of individual synaptic contacts within hippocampal neurons.

In this study, we focused on the analysis of a subset of features from our neuron cell atlases. However, the feature space can be expanded to examine any area of biological interest, such as density variations, similarities in protein distributions, or further categorization into characteristics like synaptic subtypes or more focused exploration for specific subtypes, such as phosphorylation or cellular activity. Our integrated workflow for highly multiplexed protein nano-architecture exploration can be applied to any biological system with the goal to investigate potential interactions or molecular distributions of interest. Designing a SUM-PAINT experiment requires high-quality affinity reagents for the target proteins. Given the complexity of the resulting SUM-PAINT atlas, all targets must be pre-screened, and state-of-the-art super-resolution imaging must be optimized for each target individually before transitioning to our multiplexed imaging and analysis workflow. SUM-PAINT enables hypothesis-free mapping of protein distributions and allows for unbiased feature extraction. Once interesting protein arrangements or correlations have been identified, a lower-plex experiment can be carried out, focusing on enhancing data statistics.

Despite our comprehensive analysis, we recognize that we have only begun to tap into the wealth of potential investigations afforded by the datasets we have generated, which we are excited to share with the wider scientific community. In summary, we offer an integrated data acquisition and analysis workflow with unparalleled levels of multiplexing and spatial resolution, laying the groundwork for comprehensive spatial proteomics at single-protein resolution through localization microscopy, which we term Localizomics.

### Limitations of the study

In this study, we have developed, benchmarked and applied SUM-PAINT, a method capable of virtually unlimited multiplexing for spatial profiling of molecular targets at the nanoscale. As our method relies on affinity reagents, it shares the inherent limitations of such approaches, including molecular crowding, nonspecific binding, and epitope clustering. We have devised thorough blocking procedures to minimize nonspecific binding and implemented post-fixation steps to prevent probe unbinding. However, future advancements in developing smaller, more efficient primary binders will enhance SUM-PAINT’s efficiency and practical multiplexing capabilities.

Furthermore, for the application of SUM-PAINT, we have used primary neurons, which can recapitulate several aspects of neuronal development in vitro^33, 34^, as these preparations provide optimal penetration of affinity tools and minimal background. In these cells we describe a previously unrecognized class of apparently mismatched chemical synapses that we refer to as “mixed”, containing a presynaptic vesicle pool coupled to a postsynaptic scaffold. Although we have observed similar mismatched synapses in adult mouse brain tissue, the physiological role and importance of mismatched synapses in brain development and physiology remains to be determined.

Lastly, like all localization-based super-resolution techniques, SUM-PAINT highlights the need for optical sectioning through selective plane illumination. While our current methodology allows image acquisition of single 1 µm planes, cells are several microns thick, and the elucidation of interconnections and the potential extension of this technique to tissue imaging will necessitate incorporation of advanced imaging systems such as light-sheet microscopy.

## Data and code availability

Raw localization microscopy datasets (in hdf5 format) as well as analysis scripts and software packages necessary to reproduce the results of this manuscript are available at https://www.dropbox.com/sh/woczkm7gjgbj8qx/AAD0WI7NT1TitTsnNCHHHeEra?dl=0

## Acknowledgments

We thank Isabelle Baudrexel, Alexandra Eklund, Heinrich Grabmayr and Florian Schueder for helpful discussions and technical **support**.

This research was funded in part by the European Research Council through an ERC Consolidator Grant (ReceptorPAINT, Grant agreement number 101003275), the BMBF (Project IMAGINE, FKZ: 13N15990), the Max Planck Foundation, and the Max Planck Society.

S.S. acknowledges support by the QBM graduate school. E.U., S.C.R.M. and R.K. acknowledge support by the IMPRS-LS graduate school. L.A.M. acknowledges a postdoctoral fellowship from the European Union’s Horizon 2021-2022 research and innovation program under the Marie Skłodowska-Curie grant agreement no. 101065980. M.G. acknowledges postdoctoral fellowship from the European Union’s Horizon 2020 research and innovation program under the Marie Skłodowska-Curie grant agreement no. 796606. F.O. acknowledges support by Deutsche Forschungsgemeinschaft (DFG) through the SFB1286 (project Z04). E.F.F. is funded by a Schram Stiftung (T0287/35359/2020) and a Deutsche Forschungsgemeinschaft (DFG) grant (FO 1342/1-3). E.F.F. also acknowledges the support of the Collaborative Research Center 1286 on Quantitative Synaptologie (CRC/SFB1286), Göttingen, Germany. S.S.B has received funding from F. Hoffmann-la Roche LTD (No grant number is applicable) and is supported by the Helmholtz Association under the joint research school ‘Munich School for Data Science - MUDS’. C.M. has received funding from the European Research Council (ERC) under the European Union’s Horizon 2020 research and innovation program (Grant agreement No. 866411). Part of the schematics in Figure 1 and Figure 4 were created with the help of Biorender (https://biorender.com). We thank Mark Browne (Andor Technology Ltd) for providing us with access to the Imaris image rendering software.

## Author contributions

E.M.U. developed the SUM-PAINT method, designed and conducted all DNA-PAINT and SUM-PAINT experiments, developed the spatial omics analysis workflow, analyzed and interpreted all data. S.S.B. developed the spatial omics analysis workflow, analyzed and interpreted synapse datasets. K.J. prepared neuron samples and performed antibody screening and STED imaging and analyzed data. L.A.M. analyzed and interpreted data. M.G. developed the SUM-PAINT method and performed initial experiments. Sh.S. prepared neuron samples and performed antibody screening. R.K. developed software and analyzed data. Se.S. prepared DNA-conjugated binders and performed initial experiments. S.C.M.R. developed software and analyzed data. A.P. prepared samples. C.M. supervised the spatial omics analysis workflow. F.O. supervised antibody screening, provided antibody and nanobody reagents, and interpreted data. E.F.F. supervised the neuron part of the study, prepared neuron samples, and analyzed and interpreted data. RJ conceived the concept, designed experiments, interpreted data and supervised the study. E.M.U., L.A.M., E.F.F., and R.J. wrote the manuscript with contributions from all authors. E.M.U., S.S.B., and K.J. contributed equally. All authors reviewed and approved the final manuscript.

## Methods Section

### Buffers

The following buffers were used for sample preparation and imaging:

- Buffer C+: 1× PBS, 500 mM NaCl and 0.05% Tween-20
- Buffer C: 1× PBS, 500 mM NaCl
- Antibody Incubation buffer: 1×PBS, 1 mM EDTA, 0.02% Tween-20, 0.05% NaN3, 2% BSA and 0.05 mg/ml sheared salmon sperm DNA
- Blocking Buffer: 1x PBS, 3% BSA, 0.25%Triton X-100 and 0.05 mg/ml sheared salmon sperm DNA
- Buffer A+: 10 mM Tris pH 8, 100 mM NaCl and 0.05% Tween-20
- Buffer B+: 10 mM MgCl2, 5 mM Tris-HCl pH 8, 1 mM EDTA and 0.05% Tween-20, pH 8
- Buffer B: 10 mM MgCl2, 5 mM Tris-HCl pH 8 and 1 mM EDTA, pH 8
- Optimized hybridization buffer: 10% Dextran Sulfate, 10% Ethylencarbonate, 4xSSC and 0.4 % Tween-20
- Dehybridization buffer: 10% Dextran Sulfate, 20% Ethylencarbonate, 2xSSC

### Trolox

100×Trolox was made by adding 100 mg Trolox to 430 μl of 100% methanol and 345 μl of 1 M NaOH in 3.2 ml water.

### Animals

Wild-type Wistar rat pregnant mothers or pups (*Rattus norvegicus*) and adult mice (*Mus musculus*) were obtained from the University Medical Center Göttingen and were handled according to the specifications of the University of Göttingen and of the local authority, the State of Lower Saxony (Landesamt für Verbraucherschutz, LAVES, Braunschweig, Germany). Animal experiments were approved by the local authority, the Lower Saxony State Office for Consumer Protection and Food Safety (Niedersächsisches Landesamt für Verbraucherschutz und Lebensmittelsicherheit).

### Primary cell culture

Primary hippocampal neuron cultures from embryonic day 18 (E18) Wistar rat embryos were prepared with minor adaptations from a previous work^35^. Briefly, upon dissection neurons were grown on 1 mg/ml poly-L-lysine coated coverslips over an astrocyte feeder layer and were kept in an N2-supplemented serum-free medium^34^. To prepare for the dissection of E18 rats, glial cells were prepared from P2 Wistar rat pups and seeded in 12-well plates at a density of 10000 cells per well, three days prior. Next, hippocampal neurons were seeded onto 18 mmØ coverslips at a density of 60000 cells per coverslip, with paraffin dots acting as a spacer between the neurons and glial cells. 500 µl of the cell culture medium was exchanged with fresh medium twice a week. Following this culture method, the neurons developed proper polarity, generated intricate axonal and dendritic networks, and established multiple functional synaptic connections with each other^34^. Mixed glial and neuronal cultures were prepared from P2 rats, as previously described^36^.

### Cell lines

U2OS-CRISPR-Nup96-mEGFP cells (a gift from the Ries and Ellenberg laboratories) were cultured in McCoy’s 5A medium (Thermo Fisher Scientific, 16600082) supplemented with 10% FBS. For super-resolution imaging 50k cells were seeded 24 h before fixation in glass-bottomed eight-well µ-slides (ibidi, 80827).

### Primary label design

The primary labels were shortened from 25 bases to 20 bases from the repository containing 240000 unique sequences^37^. The sequences were designed with melting temperatures between 58-68 °C. We first truncated the sequences to 20 bases by removing any five nucleotides randomly from the original sequence. We blasted the resulting sequences against the mouse genome to select the ones that show 14 bases or less homology to the genome. The remainder was further screened to contain 48% to 65% GC-content. We then evaluated each sequence both for intramolecular and intermolecular secondary structures of seven base-pairs or longer. Sequences that showed seven or more base-pair interactions were excluded from the list. The resulting sequences were selected as primary labels, extended with DNA origami staple sequences shown in Supplementary Table 1 for incorporation in rectangular DNA origami nanostructures. Secondary labels are reverse complements of the primary labels, extended with speed-optimized docking sequences^16^ at one end and an eight-nucleotide extension (GGTCTTGTGG) on the other end for toehold-mediated strand displacement.

### DNA origami sample preparation

Origami sample preparation was done in a 6-channel µ-slide (Ibidi Cat.: 80607). First 50 µl of biotin labeled bovine albumin (1 mg/ml, dissolved in buffer A+) was flushed into the chamber and incubated for 3 min. The chamber was subsequently washed with 1 ml of buffer A+ followed by incubation with 200 µl of neutravidin (0.5 mg/ml, dissolved in buffer A+) for 3 min. Afterwards the chamber was washed again with 1 ml of A+ and 1 ml of B+ buffer and incubated with biotin-labeled DNA origami (∼200 pM in buffer B+) for 3 min. Subsequently, the sample was washed with 1 ml B+ buffer and 1 ml 2xSSC buffer. Secondary label incubation was performed at a concentration of 100 nM for 15 min in optimized hybridization buffer. Finally, the chamber was washed with 5 ml of 2xSSC buffer and 1 ml of B+ buffer and 1 ml of imager solution (see Supplementary Table 3) was applied for imaging.

### U2OS Nup96-EGFP cell sample preparation

U2OS-Nup96-mEGFP cells were fixed with 4% paraformaldehyde for 20 min at room temperature. After fixation, the cells were washed three times with PBS before they were quenched using 0.1 M NH_4_Cl in PBS for 5 min. Permeabilization and blocking was performed simultaneously in Blocking buffer (3% BSA, 0.25% Triton X-100) for 45 min. Gold nanoparticles were incubated for 5 min as fiducial markers, followed by three times washing with PBS. Specific labeling of the mEGFP was done with anti-GFP nanobodies at an approximate concentration of 50 nM in Antibody incubation buffer at 4 °C overnight. The next day, the sample was washed four times with PBS and once with buffer C and 2xSSC Buffer. Secondary label incubation was performed in the optimized hybridization buffer at a concentration of 100 nM for 20 min. Afterwards, the sample was washed five times with 2xSSC buffer, once with buffer C and once with imaging solution according to supplementary table 3

### Nanobody-DNA conjugation via single cysteine

Nanobodies against GFP, tagFP, rabbit and mouse IgG were purchased from Nanotag with a single ectopic cysteine at the C-terminus for site-specific and quantitative conjugation. The conjugation to DNA-PAINT docking sites (see Supplementary Table 1) was performed as described previously20. First, buffer was exchanged to 1× PBS + 5 mM EDTA, pH 7.0 using Amicon centrifugal filters (10k MWCO) and free cysteines were reacted with 20-fold molar excess of bifunctional maleimide-DBCO linker (Sigma Aldrich, cat: 760668) for 2-3 hours on ice. Unreacted linker was removed by buffer exchange to PBS using Amicon centrifugal filters. Azide-functionalized DNA was added with 3-5 molar excess to the DBCO-nanobody and reacted overnight at 4°C. Unconjugated nanobody and free azide-DNA was removed by anion exchange using an ÄKTA Pure liquid chromatography system equipped with a Resource Q 1 ml column. Nanobody-DNA concentration was adjusted to 5 µM (in 1xPBS, 50% glycerol, 0.05% NaN3) and stored at - 20°C.

### SUM-PAINT imaging

#### SUM-PAINT DNA origami sample preparation

Origami sample preparation was done as for the secondary label origami case with the exception of DNA origami concentration being 100 pM per DNA origami, resulting in a total of 4.2 nM for all 42 different DNA origami. Secondary label incubation for barcoding was performed with 100 nM per secondary label (Supplementary Table 1), a total of 600 nM for six strands. Imager solution was applied according to Supplementary Table 3 with five times washing with buffer B in-between readout rounds. After barcoding round one, the sample was washed once with buffer B and twice with 2xSSC buffer. 100 nM per toehold strand (Supplementary Table 1), a total of 600 nM, was applied in Dehybridization buffer and incubated for 15 min. Finally, the chamber was washed five times with 2xSSC buffer and once with buffer B before proceeding to the next barcoding round (Supplementary Table 3).

#### Neuron imaging

Rat primary hippocampal neurons were fixed using 4% paraformaldehyde for 30 min at room temperature, washed four times with PBS. After fixation, neurons were quenched using 100 mM NH_4_Cl (Merck, 12125-02-9) in PBS. Then, samples were washed three times with PBS and incubated in Blocking buffer for blocking and permeabilization for 45 min. Afterwards, the samples were washed with PBS, and gold nanoparticles (1:3 dilution in PBS) were incubated for 5 min and subsequently used as fiducial markers. Primary label hybridization was performed according to Supplementary Table 3, starting with a preincubation of the antibody with their respective secondary nanobody (NanoTag Biotechnologies GmbH) in 10 µl antibody incubation buffer at room temperature for 2 h. After preincubation, an excess (molar ratio of 1:2) of unlabeled secondary nanobody was introduced for 5 min. Subsequently, a subset of six independently preincubated primary antibody and secondary nanobody complexes were pooled in 300 µl antibody incubation buffer and added to the fixed neuron sample for 60 min. Then, the sample was washed five times with PBS and once with buffer C, followed by a postfixation with 2.4 % paraformaldehyde for 7 min. Afterwards, the sample was quenched with 100 mM NH_4_Cl in PBS and finally rinsed with 2xSSC buffer. The secondary label hybridization for barcoding round 1 was then carried out according to Supplementary Table 3 with 100 nM of each secondary label for 15 min. Finally, the sample was washed five times with 2xSSC buffer and once with buffer C and imaging buffer was applied according to Supplementary Table 3. After the imaging, the sample was washed three times with 2xSSC and 600 nM (100 nM per strand) blocking strands (Supplementary Table 1) were applied to the sample for 15 min for signal extinction (optionally supplemented with 600 nM Toehold-strands). The next barcoding round containing six targets was carried out identically, with the pooling of the preformed primary antibodies and nanobody complexes (see Supplementary Table 3 for barcoding rounds of all experiments). As a last target, Actin was imaged according to Supplementary Table 3. Note that for the 12-plex demonstration (Supplementary Figure 2) primary antibody incubation was done in two subsequent steps of six primary antibodies + their respective secondary nanobodies prior to imaging.

#### Microscope setup

Imaging was carried out using an inverted microscope (Nikon Instruments, Eclipse Ti2) equipped with a Perfect Focus System using the objective-type TIRF configuration with an oil-immersion objective (Nikon Instruments, Apo SR TIRF 100X, NA 1.49, oil). A 561 nm laser (MPB Communications, 1 W) was used for excitation and was coupled into a single-mode fiber. The laser beam was passed through a cleanup filter (Chroma Technology, ZET561/10) and coupled into the microscope objective using a beam splitter (Chroma Technology, ZT561rdc). Fluorescence light was spectrally filtered with an emission filter (Chroma Technology, ET600/50m) and imaged with an sCMOS camera (Andor, Zyla 4.2 plus) without further magnification, resulting in an effective pixel size of 130 nm after 2×2 binning. The camera readout sensitivity was set to 16-bit and the readout bandwidth to 200 MHz. Image acquisition and microscope control was performed using µManager^38^. For detailed imaging parameters see Supplementary Table 3.

#### Super-resolution reconstruction

Raw DNA-PAINT data was reconstructed to super-resolution images with the Picasso software package (latest version available at https://github.com/jungmannlab/picasso). Drift correction was performed with a redundant cross-correlation following gold particles as fiducials for cellular experiments. Alignment of Exchange-PAINT and SUM-PAINT subsequent imaging rounds was performed using gold particles.

### STED imaging

#### Immunostaining of mixed synapses in primary hippocampal neurons from E18 and P2 cultures

For the time series experiment, neurons were fixed at 8, 12, 17 and 21 days *in vitro* (DIV). For the neurotransmitter receptor experiment, 22 DIV neurons were incubated live in conditioned media containing either guinea pig anti-GABA_A_ receptor γ2 subunit (1:100 dilution, Synaptic Systems, 224004) or mouse anti-GluA primary antibodies (1:100 dilution, Synaptic Systems, 182411C3, premixed with FluoTag®-X2 anti-Mouse Ig kappa light chain nanobodies conjugated to STAR580, NanoTag Biotechnologies, N1202-Ab580) for 30 minutes at 37°C. After incubation, neurons were quickly washed in cold Tyrode and fixed as described above. After fixation and quenching with 100 mM NH_4_Cl in PBS, neurons were incubated for 15 minutes at room temperature (RT) in a blocking buffer containing 2 % bovine serum albumin (AppliChem, A1391,0500) and 0.1 % Triton X-100 (Sigma Aldrich, 9036-19-5, X100-500ml) in PBS. The following steps were performed at RT. After, neurons were incubated in the same blocking buffer containing mouse anti-Gephyrin antibodies (1:200 dilution, Synaptic Systems 147011) for 1 hour. Then neurons were washed three times in PBS for 5 minutes. After primary antibody incubation, neurons were stained in blocking buffer containing secondary anti-mouse STAR635P antibodies (1:200 dilution, in-house conjugated mouse Jackson ImmunoResearch 715-005-151 and STAR635P NHS ester Abberior 07679) and anti-VGlut1 primary nanobodies conjugated to STAR580 (1:500 dilution, NanoTag Biotechnologies, N1602-Ab580-L) for 1 hour. Finally, neurons were washed three times in PBS for 5 minutes and mounted.

#### Tissue sections and immunostaining of mixed synapses in the mouse brain

To image mixed synapses in adult mouse brain tissue, mice underwent perfusion with PBS followed by 4% paraformaldehyde (PFA) in PBS. After 5 minutes of PFA perfusion, the brain was removed and further fixed overnight in 4% PFA. Coronal sections were obtained by sectioning the brain at 30 µm thickness using a vibrating microtome (VT1200S, Leica Biosystems) and stored at 4°C in PBS supplemented with 0.02% NaN_3_. For immunostaining, brain sections were first washed with PBS and quenched in 100 mM glycine (Merck, 56406) in PBS for 15 minutes. Sections were blocked in a blocking solution containing 10% Normal Donkey Serum (LIN-END9000-500, Histoprime Linaris), 1% BSA, 0.6% Triton X-100 in PBS for 2 hours. Mouse anti-Gephyrin primary antibodies were premixed at a 1 : 2.5 molar ratio with in-house produced anti-Mouse Ig kappa light chain nanobodies conjugated to STAR635P (ST635P-0003-1MG, Abberior) in 10µl for 15 minutes. Then, the premixture was diluted to 300 µl with blocking solution supplemented with anti-VGlut1 primary nanobodies conjugated to STAR580 (1:250 dilution, NanoTag Biotechnologies, N1602-Ab580-L) and tissue sections were stained with this solution overnight at 4°C. On the following day tissue sections were washed with blocking solution 3 times for 10 minutes, followed by 2 washes with high-salt PBS (PBS with 500 mM NaCl, Merck, 7647-14-5), incubated with Höchst33342 (1:10000 dilution, Thermofisher Scientific, 62249) and washed 2 times with PBS for 10 minutes. After these washes tissue sections were mounted.

#### STED sample mounting

The coverslips were quickly dipped in ddH_2_O to remove excess salts, the side of the coverslip was quickly dried on a kimwipe tissue to remove excess liquid. Immediately after, the coverslips were mounted on a microscope slide using 10 µl of Prolong Glass Antifade mounting media (ThermoFisher, P36980), left to harden overnight at RT. In the case of tissue sections, following the stainings the tissue was placed on a coverslip, the excess of PBS was cleaned with a kimwipe and mounted using Prolong Glass Antifade mounting media (ThermoFisher, P36980), which was left to harden overnight at RT. All samples were stored at 4°C until they were imaged in the following days.

#### STED image acquisition of mixed synapses

STED images were acquired using an Abberior microscope setup (Abberior Instruments GmbH) featuring an Olympus IX83 microscope body and operated with the Imspector software (version 16.3.14287-w2129 and 16.3.15521-w2209). Hippocampal CA1 *stratum radiatum* region in mouse brain tissue sections was identified by nuclear staining. Samples were imaged using a UPLSAPO100XO objective (1.4 NA), single z-plane images with a size of 20×20 µm^2^ (20×20 nm^2^ pixel size) were collected for each coverslip or a size of 60×60 µm^2^ (40×40 nm^2^ pixel size) were collected for tissue section overview. Custom size images (with a 20×20 nm pixel size) were collected in the tissue section overview region, where a mixed synapse was identified. AF488 fluorophores were excited with 485 nm laser, depleted with 595 nm pulsed laser, and the emission was detected in the range of 500-550 nm in STED mode. STAR580 and STAR635P fluorophores were excited with 561 nm and 640 nm pulsed lasers, respectively, depleted with 775 nm pulsed laser, and the emission was detected in the range of 605-625 nm and 650-720 nm, respectively, in STED mode.

#### STED image analysis of mixed synapses

Image analysis was performed using in-house written ImageJ/Fiji^39^ macro. Briefly, binary masks of VGlut1 and Gephyrin positive regions were obtained by applying a 1 sigma radius Gaussian blur filter and setting a user-defined threshold. Such objects were filtered by area (0.05 – 0.8 µm^2^ for VGlut1 and 0.02 – 0.4 µm^2^ for Gephyrin) and counted for every image. An overlap between VGlut1 and Gephyrin was manually identified by selecting a fixed size circle ROI around the overlap. Such regions were defined as “VGlut1+Gephyrin” counted and expressed as percentage over the total number of synaptic clusters (sum of VGlut1 and Gephyrin regions).

### Analysis

#### DNA origami binding kinetics and hybridization and displacement efficiency

For binding kinetics and hybridization and displacement efficiency calculation, DNA origami were handpicked using a radius of 1.2 pixels (156 nm). For the binding kinetics analysis, 400 origami were picked per dataset, for the hybridization and displacement calculation as a function of time 250 origami were picked and for the 42-plex hybridization and displacement efficiency 6440 origami were picked. Binding kinetics were then calculated on individual DNA origami sites by extracting the mean bright and dark time of all individual sites with a radius of 0.08 pixel (10.4 nm) with the picasso analysis package. Efficiency calculation was performed with single-molecule clustering, identifying the number of individual binding sites for each origami. Min sample size and cluster radius were calculated based on the localization precision of the measurement and average numbers of localizations for individual binding sites (hybridization analysis: 6 nm, 25 localizations, displacement analysis: 6 nm, 45 localization, 42-plex 5 nm, 15 localizations).

#### Synapse Segmentation

Synapse segmentation was interactively performed using the Picasso^13^ pick tool with a circular selection region of 800 nm diameter. An alignment of presynaptic and postsynaptic proteins Bassoon, Homer1 and Gephyrin displaying a clearly identifiable synaptic cleft were used as selection criteria for the first round. In a second round, the regions were selected again based on the presence of neurotransmitter transporter proteins VGlut1 and VGAT combined with common vesicle pool proteins Synaptotagmin1 and Synapsin. The selection yielded 890 synaptic regions.

#### Dataset preparation for the analysis

In this study, six datasets were analyzed. Datasets included different numbers of proteins and synapse picks, where we have Dataset 1 (21 proteins, 144 synapse picks), Dataset 2 (19, 131), Dataset 3 (17, 154), Dataset 4 (21, 186), Dataset 5 (21, 100) and Dataset 6 (14, 173) (Supplementary Figure 5, Supplementary Table 3)

#### Feature extraction

The feature extraction process was divided into (i) histogram feature extraction and (ii) clustering feature extraction. For histogram feature extraction, a 3D histogram of the super-resolution localizations of the proteins was calculated. For each histogram, the localizations were decentred and normalized by mapping to [-1,1]^3^. Then the 3D histogram with 8000 3D bins (20 bins in each dimension) was calculated. This histogram generation allowed the calculation of, mean, standard deviation, skewness, kurtosis, entropy, minimum, maximum and entropy (all unitless due to normalization). In addition, the pairwise Wasserstein distance^40^, Jensen Shannon distance^41^, and cosine distance of the histograms of different proteins were obtained. For clustering feature extraction, spatial clustering using DBSCAN^42^ was performed for each protein species in each segmented synapse. DBSCAN parameters, minimum localizations, and clustering radius were selected based on the imaging parameters of the individual super-resolution channel of the protein. Background regions were taken as a base for determining the cluster minimum localization number and both DBSCAN parameters were further adjusted based on visual validation on selected synapses to determine a cutoff value distinguishing background from specific protein clusters. All clustering parameters for the individual datasets and protein channels can be found in Supplementary Table 5. Afterward, the largest cluster based on the number of localizations for each protein was classified as the main cluster. In addition to the number of localizations, spatial cluster size, Center of Mass (CoM) and convex hull volume (unit=px^3^) of each cluster were calculated. In addition, we used α-shape^43^ with α = 0.5 to calculate the cluster volume (unit=px^3^) and surface area (unit=px^2^). Finally, the distance between the CoM (unit=px) of each protein species was calculated (Supplementary Table 4). Considering that some of the datasets did not contain all of the proteins, the features were input with 0 to enable further numerical evaluations. For the analysis of single features with pixel-based units, these features were transformed to nm-based values to provide meaningful values.

In total, 2537 features were extracted for each of the 890 synapses. This feature space contained 171 histogram-based features (HF) for a single protein channel and 1450 pairwise histogram-based features (HD), comparing features between protein targets. On the spatial clustering feature side, our extraction contained 796 clustering-based features (CF) for single-protein channels and 48 features comparing the CoM distance between protein species (CD). Finally, for the α-shape based volume and surface area, we extracted 72 features for single-protein channels (AS). To assemble a comparable feature selection between the six datasets, only proteins imaged in all six datasets were considered for the analysis Moreover, constant features (i.e. no variance) were excluded from the analysis as they provided no information, yielding to 1590 features.

#### Dimensionality reduction and clustering

To reduce the dimensionality of the datasets to aid data interpretation, we used Uniform Manifold Approximation and Projection (UMAP). Since we were pooling segmented synapses from different datasets, the slight variation of experimental conditions was due to minor changes in sample preparation or different imaging parameters. We hence used feature normalization to be able to compare synapses from different datasets robustly. For an extracted feature, *f* we calculated 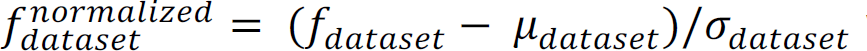 where µ*_dataset_* and σ*_dataset_* are mean and standard deviation of feature *f* among all synapse picks per dataset. For example, when we consider the number of localizations as a single feature *f_dataset_*, there are variations between the datasets because of different imager concentrations and positions of the cell in the imaging volume. We then calculate 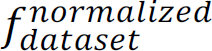 considering µ*_dataset_* and σ*_dataset_* of all synapses in the datasets, yielding features with mean=0 and standard deviation=1 for each dataset. Therefore, the features among the datasets can be comparable after the normalization.

To reduce the dataset’s high dimensionality while learning the feature space’s structure, uniform manifold approximation and project (UMAP) was used^26^. The reason for this choice is UMAP’s robustness in dealing with high-dimensional data and its efficiency in terms of run time^44^. We used the Python implementation of umap-learn (version=0.5.1) with all its default parameters, including n_neighbors=15, min_dist=0.1, n_components=2, and metric=’euclidean’. This reduced 1590 features to a 2-dimensional space (n_component = 2). Next, we focused on understanding the structure of the 2-dimensional space. Therefore, we applied KMeans clustering^45^ with k = 3 from scikit-learn (version=0.24.2) with its default parameters. The resulting clusters from the UMAP, Figure 5B were then assigned as the three synaptic subtypes under investigation, namely inhibitory, mixed and excitatory synapses. We also tried spectral clustering which consistently yielded 3 clusters, but preferred KMeans as it provided a better separation among the clusters. To visualize the variation of a single feature (e.g. numbers of localizations of Bassoon) on an individual synapse level we rendered the UMAP with a grayscale color for each synapse ranging from minimum value (white) to maximum value (black) for the given feature.

#### Center of Mass Matrix

The CoM distance matrix was assembled based on the 48 features extracted as pairwise CoM distances for protein clusters. The single pairwise CoM between two protein species is the absolute distance between the CoM of DBSCAN-detected protein clusters in a single synapse. To assemble the matrix, only synapses that expressed a spatial cluster in both protein channels were taken into account. For the overall CoM matrix (Figure 6C), the average of all segmented synapses was taken to calculate the average CoM distance, while the synapse subtype-specific proteins Homer1, Gephyrin were pooled as a postsynaptic scaffold (PSS), and VGlut1 and VGAT were pooled as neurotransmitter transporter (NTT). For the assembly of synapse subtype-specific CoM matrices (Supplementary Figure 6), the synapse subtype-specific proteins were separated.

#### Relative abundance analysis and spatial Cluster Volume

For calculating the relative abundance of a given protein species among the three synapse subtypes, we considered the detected and not detected clusters by spatial clustering with DBSCAN. If the segmented synapse is expressing a protein cluster, which was detected by DBSCAN we label this protein species with a 1. If the cluster detection threshold is not sufficient, we assign a 0. The relative abundance is then calculated as the ratio between the number of detected spatial clusters for a given protein and the total number of synapses. As an example, since the synapse selection was based on a positive DBSCAN spatial clustering on Bassoon, all synapses express a protein cluster in this protein channel and the relative abundance for Bassoon is 100%. To further investigate the specifics of the spatial protein clusters detected by DBSCAN, we plotted the average absolute volume calculated by α-shape in a heatmap, separated for the synaptic subtypes.

#### Distance Profile

To assemble a representative distance profile for each synaptic subtype, three synapses, one for each subtype, were selected and rotated *en-phase*. We then plotted the localizations for each protein channel of the three synapses in a violin plot, with the distance between the mean of the pre- and postsynaptic scaffold proteins as 0. The polarity of the synapse was assigned such that the postsynaptic scaffold would be on the negative side and the presynaptic scaffold on the positive. Only protein channels in which the DBSCAN spatial clustering detected a relevant cluster for the selected synapse were taken for representation.

#### Volume ratio correlation

To make a conclusive comparison between the expression level of different proteins, we considered the volume of a given protein species, calculated by α-shape as a proxy. The volume of the protein is the only feature that can be considered for abundance comparison between protein species since the number of localizations depends heavily on the labeling efficiency of the antibody. We first separated the synapses based on their subtype, then plotted the pairwise volume ratio of each protein rendering each individual synapse and calculated the Pearson correlation for the two proteins. For example, we can consider the volume correlation between the synaptic vesicle protein Synaptotagmin1 and the NTT. Since both proteins are associated with the vesicle pool, the direct comparison between the volume ratio of those proteins yields relatively high correlation values for all three synaptic subtypes (r = 0.82 for excitatory, r = 0.6 for inhibitory, r = 0.86 for mixed). We assembled an 11x11 matrix showing all correlation values (grayscale 0 to 1) for an overall comparison of all pairwise protein values. To compare these matrices between the synaptic subtypes, we kept the protein order the same for each matrix, allowing a direct visual inspection of differences and similarities. Finally, to make a more quantitative comparison of the volume correlation within one synaptic subtype, we performed hierarchical clustering and assembled a dendrogram for each matrix. The hierarchical clustering was based on the nearest neighbor point algorithm and Euclidean distance^46^ based on SciPy (version=1.8.0) implementation.

#### Statistical tests among features and clusters

To assess if there is a significant difference between the distributions of synapse volumes and the number of localizations of individual protein species among the three synaptic subtypes in Figure 6 H-J we systematically compared the features with Wilcoxon-Mann-Withney U Test^47^. This test is a non-parametric test with no assumption on the distribution of the data. A significant difference between distributions in this test is indicated by a p-value < 0.05, indicated by an asterisk (*) in the figures. To correct for multiple testing and reduce the false discovery rate^48^, all p-values were corrected using the Benjamini-Hochberg procedure^49^.

## Supplementary Figures

**Supplementary Figure 1.**
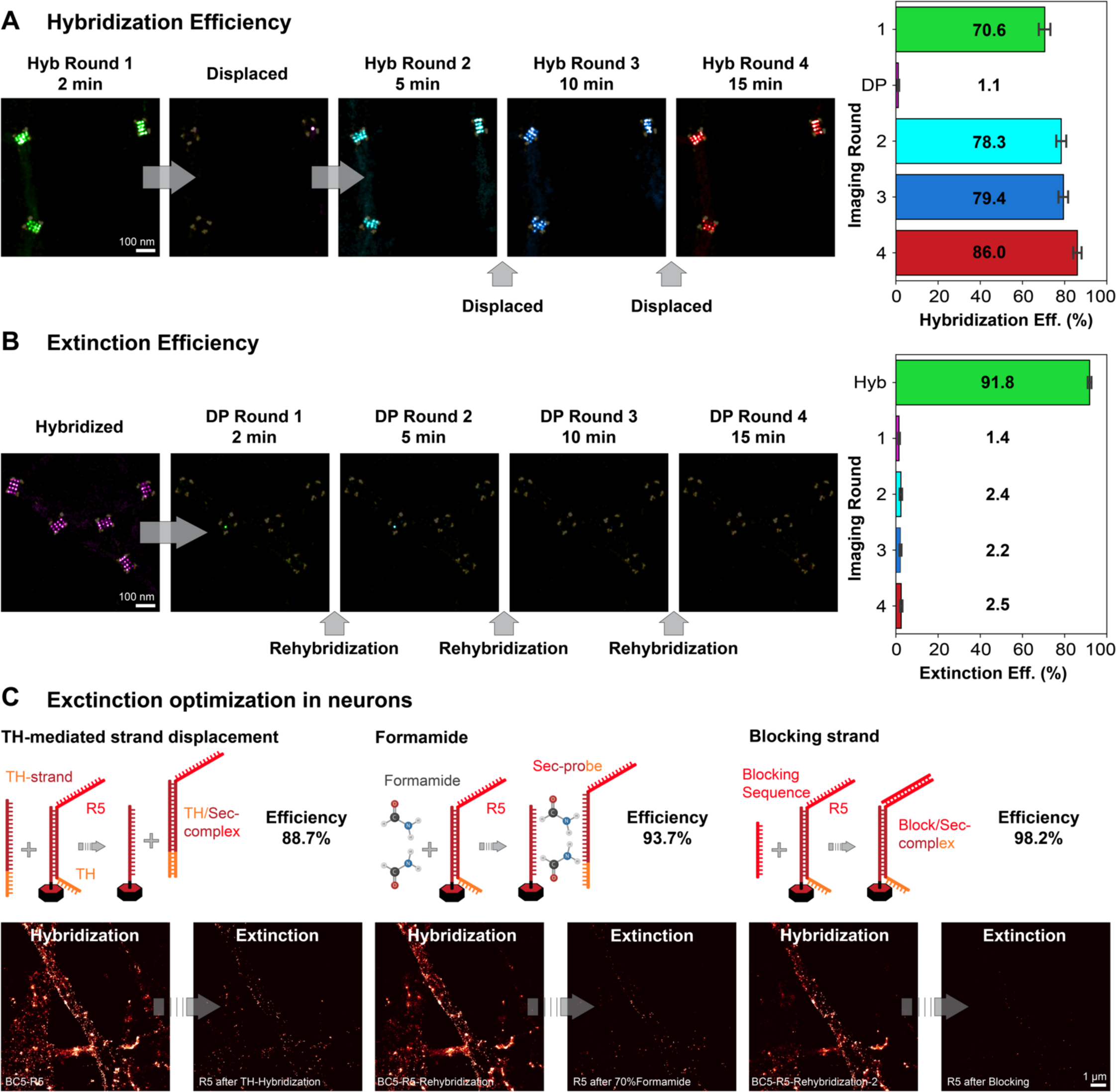
Signal hybridization and extinction optimization. **(A)** Hybridization efficiency optimization on DNA origami. The hybridization efficiency is evaluated by a time-series featuring 2 min, 5 min, 10 min and 15 min of hybridization time, each followed by 15 min of signal extinction by toehold-mediated strand displacement. The resulting labeling efficiency (a combination of primary barcode incorporation and secondary label hybridization efficiency) of 250 origami and one exemplary experiment is shown by a bar plot. **(B)** Toehold-mediated strand displacement efficiency optimization on DNA origami. The Displacement efficiency is evaluated by a time-series featuring 2 min, 5 min, 10 min and 15 min of displacement time, each followed by 15 min of signal rehybridization. The resulting extinction efficiency of 250 origami and one exemplary experiment is shown by a bar plot. **(C)** Signal extinction optimization in neurons. Extinction in neurons is evaluated using three different methods, signal removal by toehold(TH)-mediated strand displacement, signal removal by formamide denaturing agent and signal blocking by blocking strand hybridization. The extinction efficiency is calculated as the remaining signal in the exemplary field of view compared to the hybridized or re-hybridized signal.

**Supplementary Figure 2.**
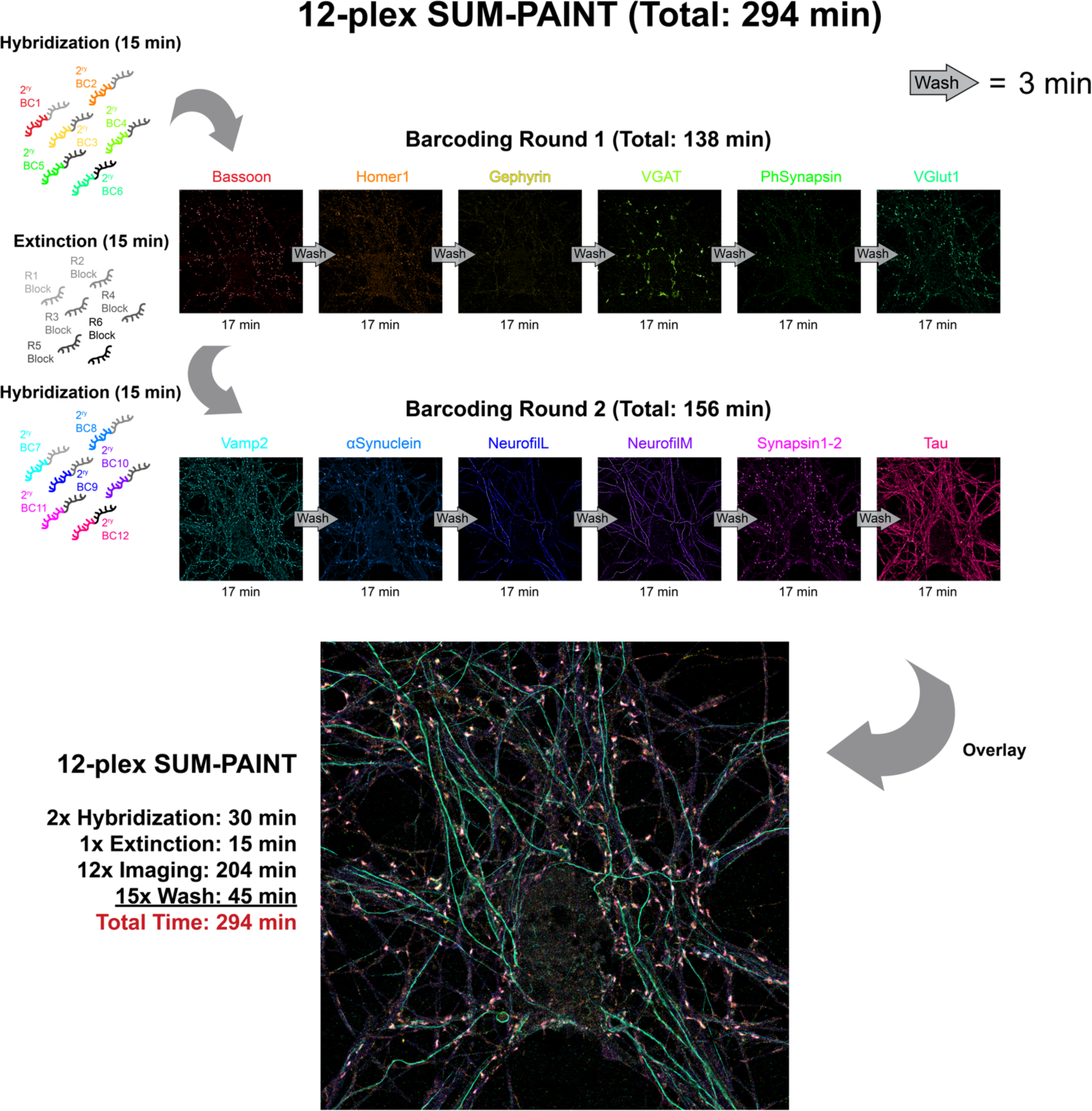
12-plex SUM-PAINT demonstration. Exemplary 12-plex SUM-PAINT imaging workflow and result. 12-plex SUM-PAINT imaging can be completed in roughly 5 hours. Hybridization time is 15 min followed by 6x17 min of super-resolved imaging with R1 to R6 sequences in barcoding round 1. Subsequent signal extinction is done by blocking strand hybridization (15 min) and followed by the hybridization of the second-round labels. Barcoding round 2 is then carried out in the same way as round one (6x17 min) resulting in a total experiment time of 294 min. Images show the individual protein targets and their composite 12-plex overlay.

**Supplementary Figure 3.**
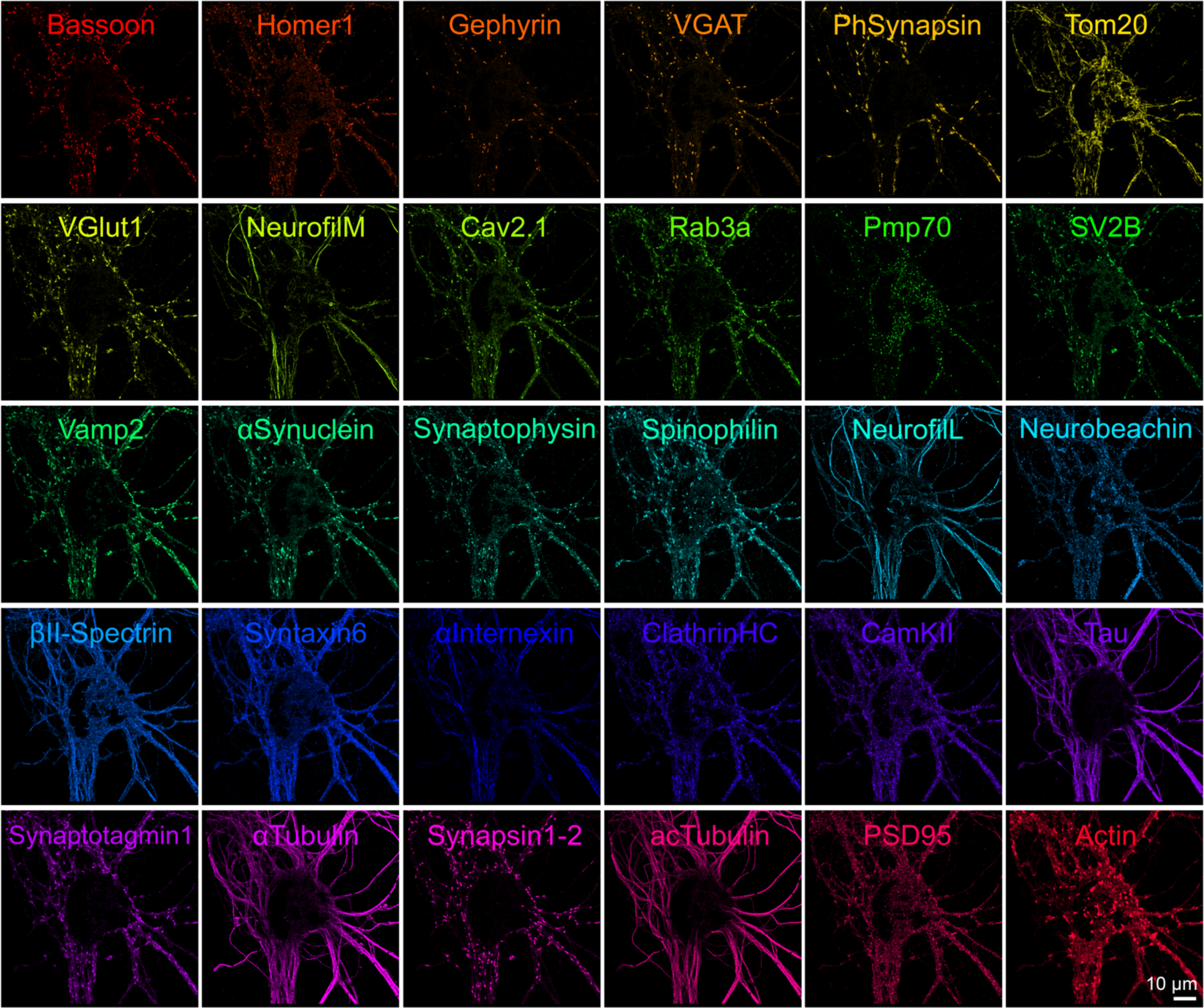
Gallery of individual protein targets of the 30-plex SUM-PAINT experiment. DNA-PAINT imaging rounds were completed in 17 min for each protein target yielding an average localization precision of 6.6 nm. The protein targets range from synaptic scaffold proteins (Bassoon, Homer1, Gephyrin, PSD95) to vesicle pool proteins (e.g. VGlut1, VGAT, Vamp2, Synaptotagmin1) to organelle markers (Tom20, Pmp70) and to cytoskeleton proteins (e.g. βII-Spectrin, ac-Tubulin, NeurofilamentM/L, Actin).

**Supplementary Figure 4.**
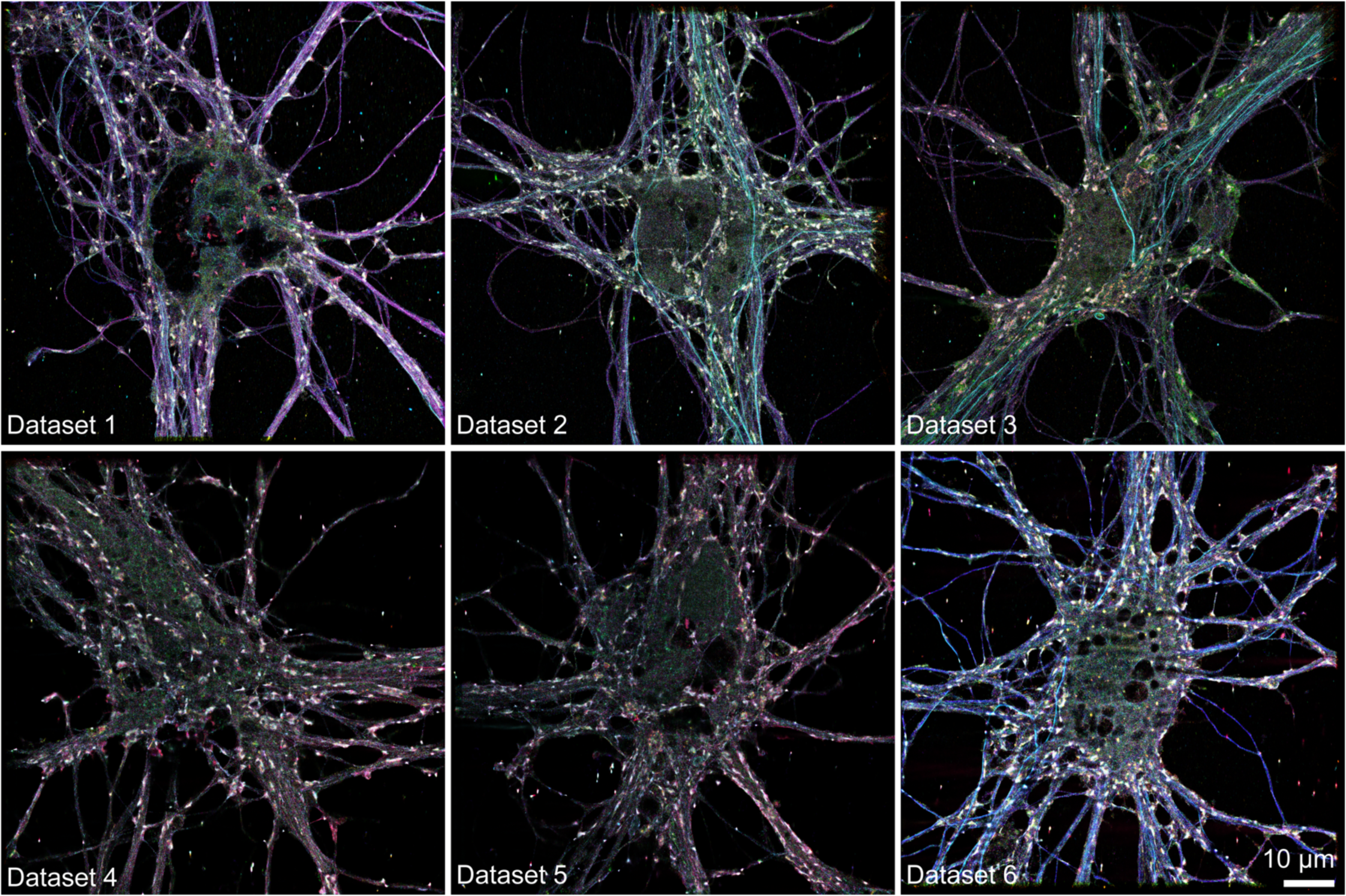
Six multiplexed neuron atlas datasets. Each dataset was imaged with at least 19 different protein targets (see Supplementary Table 3).

**Supplementary Figure 5.**
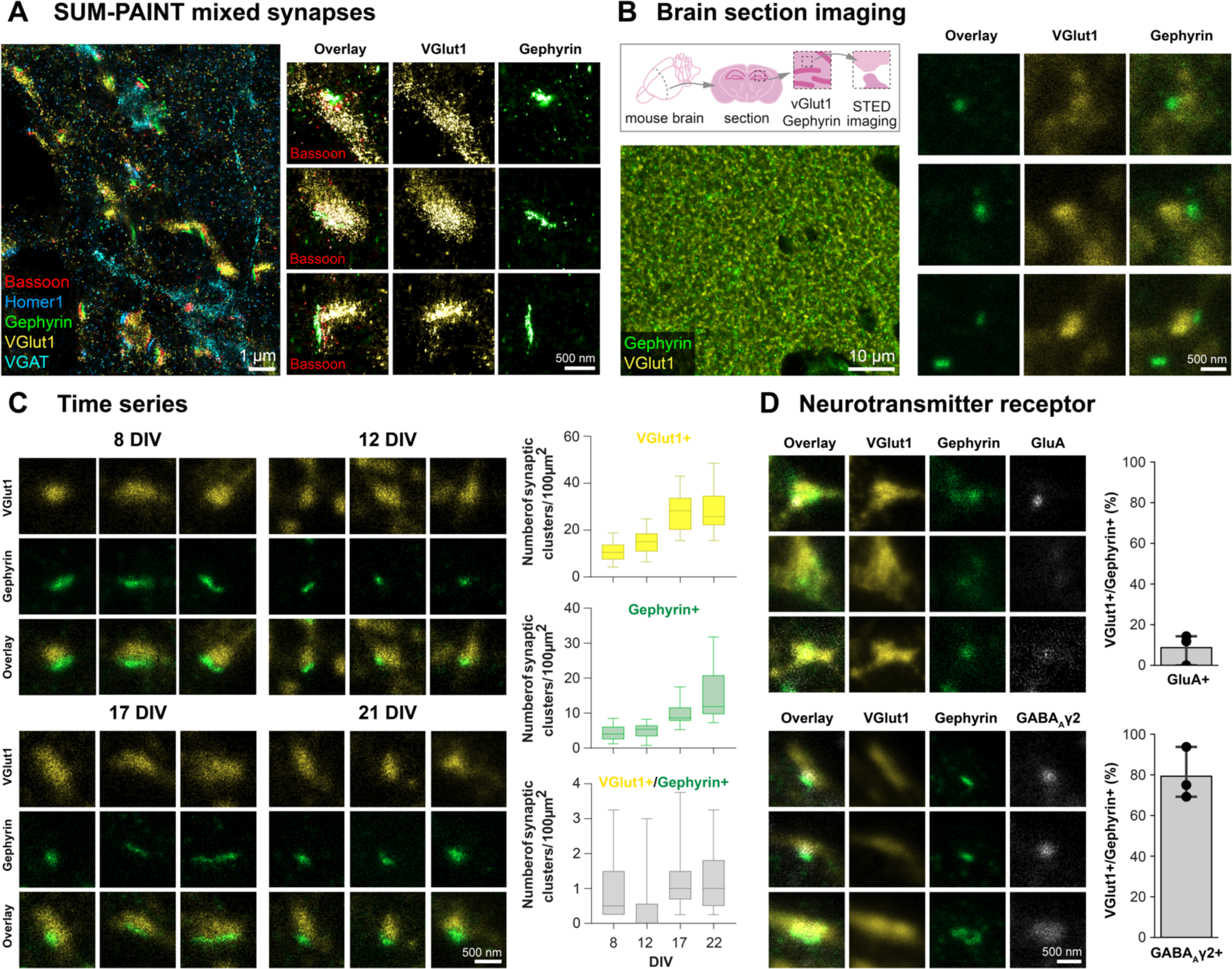
Characterization of the mixed synapse. **(A)** 5-plex super-resolved overlay of synaptic scaffold proteins Bassoon, Homer1 and Gephyrin with NTT markers VGlut1 and VGAT next to three exemplary mixed synapses. Mixed synapses are shown with separated Gephyrin (green) and VGlut1 (yellow) channels and an overlay with Bassoon (red) in addition. **(B)** Brain section imaging with 2-color STED microscopy. The upper left scheme indicates the location of the imaged region in the mouse hippocampus (*stratum radiatum*). The bottom left panel shows a 2-color overlay of an exemplary hippocampal region with Gephyrin (green) and VGlut1 (yellow). The right panels show three exemplary mixed synapses with Gephyrin and VGlut1 both as single channels and in a merged overlay. **(C)** Time series evaluating the possible developmental effect on the abundance of the mixed synapse type revealed with STED microscopy. DIV 7, 12, 15 and 21 neurons were imaged with 2-color STED microscopy with three representative mixed synapses for each time-step. The overall evaluation of synaptic puncta is shown in three box-plots for VGlut1^+^, Gephyrin^+^ and VGlut1^+^/Gephyrin^+^ puncta, indicating that overall synaptic puncta numbers rise until DIV 17, while the mixed class stays at a constant low abundance of about one synapse per 100 µm^2^. **(D)** Postsynaptic neurotransmitter receptor characterization for the mixed synapse type. 3-color STED imaging with VGlut1 (yellow), Gephyrin (green) and the postsynaptic receptors GluA1 (gray) or GABAAγ2 (gray) with three exemplary mixed synapses for each experiment. The evaluation shows that ∼9 % of the mixed synapses express GluA as a postsynaptic receptor, while ∼79 % express GABAAγ-2.

**Supplementary Figure 6.**
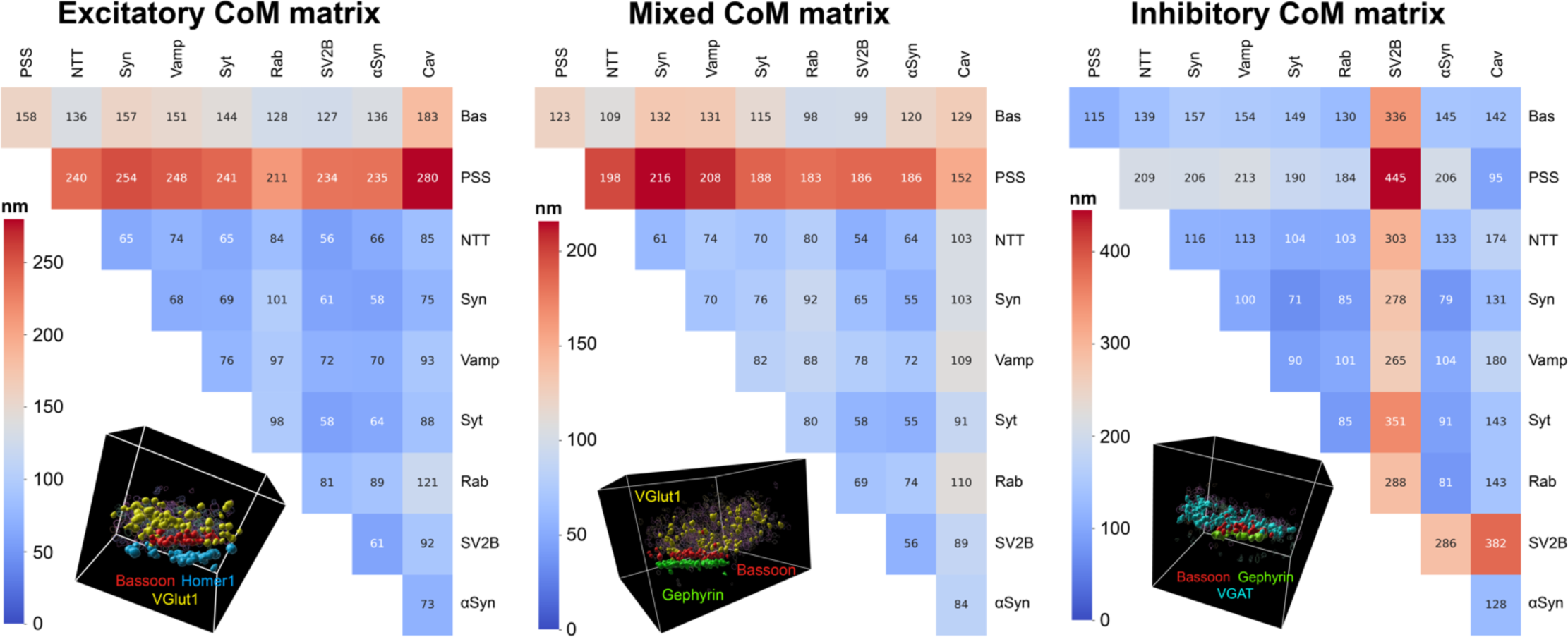
Center-of-Mass distances for each synapse subtype. Average Center-of-Mass (CoM) distance matrices for excitatory, inhibitory and mixed synapses, respectively. The distances for the protein clusters CoM were calculated for each synapse individually and the average of all excitatory (411), inhibitory (189) and mixed (297) synapses were subsequently assembled into a 9x9 matrix.

